# Functional and molecular profiles of the neuroretina and retinal pigmented epithelium-choroid in response to aging and Nrf2 deletion

**DOI:** 10.64898/2026.10.08.757667

**Authors:** Becca L. Wells, Adrian Filiberti, David R. Stanford, Kevin D. Pham, Dorothy A. Walton, Scott M. Plafker, Willard M. Freeman, Ana J. Chucair-Elliott

## Abstract

**Purpose:** Age is the leading factor for age-related macular degeneration (AMD), in which retinal pigmented epithelium (RPE) atrophy is a hallmark. The nuclear factor-erythroid 2-related factor 2 (Nrf2), a major anti-stress transcription factor, decreases in AMD, suggesting bioenergetic stress interacts with age during AMD pathogenesis. We profiled visual function and gene expression of the retina and RPE-choroid with Nrf2 deletion and aging.

**Methods:** We tested 3mo and 14mo Nrf2 null (Nrf2 KO) and Nrf2 WT mice for optokinetic tracking (OKT) and dark-adapted electroretinography (ERG) responses. Eyes were harvested for retina outer nuclear layer (ONL) thickness analysis and retina and RPE-choroid cup RNA isolation for RNAseq profiling.

**Results:** Visual acuity was reduced with aging with lower values in the Nrf2 KO vs Nrf2 WT comparison at 14mo. We observed aged-related reduction of a-, b-, and c-wave ERG amplitudes in both WT and KO mice. Age-related c-wave decline was exacerbated by Nrf2 loss. By RNAseq, two-way ANOVA of RPE-choroid samples revealed differentially expressed genes (DEGs) with age and genotype main effects, as well as DEGs with age and genotype interaction effects. Age DEGs were over-represented in pathways related to inflammation and phagocytosis. Genotype DEGs were associated with alterations in antioxidant and inflammatory processes. Retina RNAseq showed age and genotype DEGs.

**Conclusions:** Ablation of Nrf2 exacerbates age-related declines suggesting that decrease in Nrf2 with aging has functional consequences for inflammation/resiliency. We provide retina and RPE-choroid bulk-RNAseq datasets from the same cohort of mice that can be interrogated to better understand degeneration of retina/RPE-choroid.

## Introduction

Aging is the greatest non-modifiable risk factor for age-related macular degeneration (AMD), a multifactorial blinding disease in people over 60 years old^1,2^. Environmental factors known to increase oxidative stress, such as diet, obesity, and smoking, are also linked to higher risk of developing AMD^3,4^. A variety of ocular and systemic cells have been implicated in the development of the disease, but ample evidence supports that the choroid and retinal pigmented epithelium (RPE) are important sites of AMD pathogenesis^5–9^ which leads to photoreceptor cell loss.

The retina is a highly metabolic tissue with high oxygen demand, delivered by a combination of choroidal vascular bed and retinal vasculature^10^, which relies on the trophic, metabolic, and functional support of the RPE^11^. RPE cells are located between photoreceptor outer segments and Bruch’s membrane, adjacent to the choriocapillaris layer of the choroid. Through its functional interactions with retina and underlying choroid, the RPE plays critical roles in maintaining health of both tissues (reviewed in ^11–14^). However, exposure to excessive oxidative stress can cause cellular damage in various ocular cell types that disrupt homeostasis and contribute to disease onset^15^.

Activation of protective pathways and upregulation of endogenous antioxidants are cellular responses against excessive oxidative stress. In a wide range of tissues, Nrf2 (*Nfe2l2*) is the key transcription factor for sensing and neutralizing oxidative stress^16^. Nrf2 induces the expression of antioxidant and cytoprotective enzymes via binding to the cis-acting antioxidant response element (ARE) in the promoter of its target genes. In addition, Nrf2 regulates pathways that generate NADPH, which in turn regenerates restored oxidized antioxidants to their reduced forms^17,18^ to sustain mitochondrial bioenergetics^19^, fatty acid oxidation and ATP synthesis^19,20^. Apart from regulating the cellular redox status and bioenergetics, Nrf2 acts as a negative regulator of inflammation^21–24^. Given the high oxygen tensions needed to match the metabolic demands, the Nrf2-regulated antioxidant defense system is especially critical in both retina and RPE. As such, Nrf2 is expressed in retinal cell types such as Müller glial cells (MGC) and ganglion cells (RGCs), and in the RPE^25–27^.

Of clinical relevance, Nrf2 expression in the RPE has been shown to be altered with chronic oxidative stress such as with early AMD^28^. In line with these observations, and despite that mice lack a macula, several preclinical studies support the importance of Nrf2 signaling for retina/RPE, suggesting relevance to human disease. For instance, loss of Nrf2 rendered mice susceptible to vascular damage by ischemia-reperfusion injury^29^. Ablation of Nrf2 also exacerbated visual deficits, inflammation of the optic nerve, and RGC degeneration in a murine model of multiple sclerosis^30^. Nrf2 deficient mice displayed higher susceptibility to chemically-induced oxidative stress with aging^31^, as well as to the development of retinopathy, including AMD features, between 12-16 months of age^32,33^. In addition, in response to a high glycemic index diet, Nrf2 null mice developed atrophic AMD-like disease that was prevented by a low glycemic diet^34^. Further, Nrf2 over-expression specifically in RPE elicited functional and structural improvements when tested in mouse models of retinal degeneration^35^. While these studies have provided information on the role of oxidative stress in disease pathogenesis, how deficiency of Nrf2 intersects with normal aging, the main risk factor for AMD, to affect gene expression programs and function of the retina and RPE remains unresolved. While Nrf2 deficient mice are often used to model relevant aspects of retinal disease and to test neuroprotective modalities, no transcriptomic datasets comparing Nrf2 WT and Nrf2 null (Nrf2 KO) mouse retina and RPE/choroid with age are publicly available, which prompted us to fill this data gap in the field.

Here, we profile visual function and bulk-level transcriptomes from female and male mouse retina and RPE-choroid tissues at 3 months and 14 months of age. We report an interaction effect of aging and Nrf2 expression status to reduce visual acuity and electroretinographic c-wave response, consistent with differential age and genotype effects on RPE-choroid gene expression that reveal alterations in inflammatory pathways with age in the absence of Nrf2. As expected in the absence of injury stimulus, these alterations did not involve frank changes in expression of genes coding for the antioxidant system. Of interest, our analyses support that with aging and higher susceptibility to oxidative stress, RPE and choroidal impairment precedes retina functional/structural decline. The data provided here can be used to better understand mechanisms driving degeneration of retina/RPE with potential clinical implications since aging and oxidative stress are risk factors for AMD.

## Methods

### Animals

All animal procedures were approved by the Institutional Animal Care and Use Committee at the Oklahoma Medical Research Foundation (OMRF) and adhere to the ARVO Statement for the Use of Animals on Ophthalmic and Vision Research. Nrf2^−/−^ (Nrf2 KO) mice^32,36^, originally generated in a 129SVJ background, were purchased from the Jackson Laboratory (stock number # 017009, Bar Harbor, ME) and backcrossed into C57BL/6J for greater than ten generations under SPF conditions in a HEPA barrier environment^30^. Nrf2^+/+^ mice generated by backcrossing were used as WT controls. Mouse colonies were housed and maintained in the OMRF vivarium until functional testing and tissue harvesting. Mice (males and females) were used at 3 and 14 months old at the time of performing experiments as detailed below. DNA was extracted from mouse ear punch samples for genotyping. All experimental mice were screened for Pde6b^rd^^1^ (*Rd1)* ^37^ and Crb1^rd8^ *(Rd8)*^38^ mutations. All mice were *Rd1^wt/wt^* and *Rd8^wt/wt^*. Euthanasia prior to tissue harvesting was carried out by cervical dislocation and decapitation, in accordance with AVMA guidelines ^39,40^.

### Visual acuity assessment

Visual acuity threshold was measured with the optokinetic tracking (OKT) response using Optometry software and apparatus (Cerebral Mechanics Inc., Alberta, Canada) as previously described^41^. Briefly, mice were surrounded by a virtual cylinder consisting of vertical lines rotating at varying frequencies, and tracking behavior was assessed by an investigator blinded to genotype and age, as previously reported^30,42^. Visual acuity is represented as the highest spatial frequency at which mice track the rotating cylinder. Because tracking is a temporal to nasal-specific reflex, counter-clockwise and clockwise rotations exclusively test the right eye and the left eye, respectively.

### Electroretinography (ERG)

ERG recordings were performed as previously described^43^, from 10:00 am to 2:00 pm ^44,45^. After overnight dark adaptation, mice were anesthetized with an intraperitoneal injection of ketamine (100 mg/kg) and xylazine (10 mg/kg), placed on a heating pad at 37°C, and administered a drop of 0.5% proparacaine onto their corneas. Mouse pupils were then dilated topically with sequential drops of 0.5% tropicamide and 10% phenylephrine. After 10 min of topical drug administration, anesthetized mice were placed on a controlled heater for the duration of the recordings. A dual function stimulator-electrode probe was placed directly onto the corneal surface, after topical application of 1% methylcellulose, to simultaneously deliver light stimuli and record electrical activity. Full-field dark-adapted (a- and b-waves) and c-wave ERGs were recorded from both eyes using a Celeris Next generation rodent ERG testing system (Diagnosys LLC, Lowell, MA)^43,46^. With this system, stimuli were delivered to one eye at a time, with the fellow, unstimulated eye, serving as reference. For full-field scotopic ERG recordings, a series of increasing flash intensities (0.01, 0.1, and 1 cd.s/m^2^) was presented. For flash responses, the a- and b-wave amplitudes and implicit times (T_a_ and T_b_) were calculated according to convention at the peaks and troughs^47^. RPE-driven c-wave responses were elicited by a 0.9 cd.s/m^2^ intensity flash and recorded within a period of 10 s after the flash presentation.

### Tissue harvesting

Following ERG recordings, mice were euthanized and one eye was collected for histology purposes while the fellow eye was dissected out to collect separately the retina and eyecup (RPE-choroid) for RNA extraction and RNAseq.

### Histology, imaging, and retina morphometry

After ERG recordings, mice were euthanized and the eyes were enucleated and fixed in Prefer (glyoxal, ethanol, buffer) (Anatech Ltd.) for 2-4 h and placed in 70% ethanol at room temperature until embedded in paraffin. Sagittal sections through the center of the eye, including the optic disc, were cut at 5-μm thickness (Excalibur Pathology, Moore OK). Hematoxylin and eosin staining was performed to examine eye morphology and capture whole section images using a Motic EasyScan One Slide Scanner (Motic Instruments, Inc., San Francisco, CA) at 40X at the Dean McGee Eye Institute-OUHC. All images were acquired using identical scanning parameters and illumination settings across all samples. From captured images, the thickness of the outer nuclear layer (ONL) was measured at consecutive positions spaced 0.2 mm apart from the optic nerve (ON) to the inferior and superior ora serrata. Measurements were performed using Fiji/ImageJ (NIH) with calibrated spatial scaling. Due to a lack of marking of the superior aspect of the collected eyeballs, we were agnostic to superior/inferior identities of retina hemispheres. Thus, the inferior and superior ONL thickness values measured at equidistant points from the ON were averaged and plotted as a “hemi”-spidergraph.

### RNA extraction

Dissected retina and eyecup (RPE-choroid) samples were collected from euthanized mice immediately following ERG recordings. Eyecups and retinas from all experimental groups were individually placed in 1.5 ml tubes, snap frozen, and stored at −80 °C until processing. Each *retina sample* was homogenized in 150 µl ice-cold homogenization buffer (50 mM Tris, pH 7.4; 12 mM MgCl2; 100 mM KCl; 1% NP-40; 1 mg/ml sodium heparin; 1 mM DTT) supplemented with 100 µg/ml cycloheximide (#C4859-1ML, Millipore Sigma), 0.5 mM spermidine (#05292-1ML-F, Millipore Sigma), 200 units/ml RNaseOUT Recombinant Ribonuclease Inhibitor (#10777019; Thermo Fisher), and 1X cOmplete, EDTA-free Protease Inhibitor Cocktail (#11836170001; Millipore Sigma), using pellet pestle cordless motor and blue pestle with two pulses of 10 sec/each (on ice). Retina homogenates were taken up to 500 µl volume with homogenization buffer. After centrifugation (12,000 x g, 10 min, at 4° C), 100 µl of the supernatant were mixed by pipetting with 350 µl of Buffer RLT (Qiagen) supplemented with 2-β mercaptoethanol (RLT/BME) and subsequently mixed with 350 µl ethanol. The entire final volume was loaded onto a RNeasy MinElute column and RNA was isolated using RNeasy Mini Kit (#74104, Qiagen), according to manufacturer’s instructions, and including a DNAse I treatment (#79256, Qiagen). Each *eyecup sample* was homogenized in 150 µl RLT/BME using pellet pestle cordless motor and blue pestle with two pulses of 10 sec/each (on ice). After taking up to 350 µl final volume with RLT/BME and centrifuging (18,000 x g for 3 minutes at 4°*C*), the supernatant was mixed with 350 µl ethanol and the entire final volume was loaded onto an RNeasy MinElute column for RNA extraction as described for retina samples. Retina and eyecup RNA were quantified with a Nanodrop One spectrophotometer (Thermo Fisher Scientific) and nucleic acid quality assessed by HSRNA ScreenTape with a 4150 TapeStation analyzer (Agilent Technologies, Santa Clara, CA).

### Library construction and RNA sequencing (RNAseq)

The NEBNext Ultra II Directional Library Prep Kit for Illumina (#NEBE7760L; New England Biolabs Inc., Ipswich, MA) was used with 100 ng of total RNA for the preparation of strand-specific sequencing libraries from retina- and eyecup-isolated RNA samples according to manufacturer’s instructions^40^. Briefly, polyA containing mRNA was purified using oligo-dT attached magnetic beads. mRNA was chemically fragmented and cDNA synthesized. For strand-specificity, the incorporation of dUTP instead of dTTP in the second strand cDNA synthesis does not allow amplification past this dUTP with the polymerase. Following cDNA synthesis, each product underwent end repair process, A tailing and adapter ligation. The cDNA products were further purified and enriched using PCR to make the final library for sequencing. Library sizing was performed with HS DNA ScreenTape (#5067-5584; Agilent Technologies). The libraries for each sample were pooled at 4 nM concentration and sequenced using an Illumina NovaSeq X Plus system (S4 PE150) at the OMRF Clinical Genomics Facility.

### RNAseq Data Analysis

Following sequencing, reads were trimmed, aligned, differential expression statistics, and correlation analyses were performed using Strand NGS software (Version 4.0, Bangalore, India)^40,48^. Reads were aligned against the Mm39 build of the mouse genome with RefSeq Genes and Transcripts (2024.03.14) annotation build. Alignment and filtering criteria included: adapter trimming, fixed 2bp trim from 5’ and 6bp from 3’ ends, a maximum number of one novel splice allowed per read, a minimum of 90% identity with the reference sequence, a maximum of 5% gap, trimming of 3’ end with Q<30. Alignment was performed directionally with Read 1 aligned in reverse and Read 2 in forward orientation. Reads were filtered based on the mapping status and only those reads that aligned normally (in the appropriate direction) were retained. Duplicate aligning reads were then removed. Normalization was performed with the DESeq algorithm. Transcripts with an average read count value ≥20 in at least 100% of the samples in at least one group were considered expressed at a level sufficient for quantitation and those transcripts below this level were considered not detected/not expressed and excluded, as these low levels of reads are close to background and are highly variable. For those transcripts meeting this statistical criterion, a fold change >|1.25| cutoff was used to eliminate those genes which were statistically significant but unlikely to be biologically significant and orthogonally confirmable due to their very small magnitude of change. For statistical multiple comparison analysis of differential expression, 2-way ANOVA with age and genotype or age and sex main effects with Benjamini-Hochberg multiple testing correction was performed (fold change >|1.25| cutoff, corrected p value ≤0.05). For genes showing significant age or genotype effects, differences among individual age/genotype groups were evaluated by 1-way ANOVA with Student Newman Keuls (SNK) pairwise post-hoc testing with Benjamini-Hochberg correction. Visualizations of hierarchical clustering and principal component analysis (PCA) were performed in Strand NGS. The entirety of the sequencing data discussed in this manuscript is available for download in FASTQ format. Data have been deposited in NCBI’s Gene Expression Omnibus from NCBI Sequence Read Archive^49^. Lists of differentially expressed genes were imported into the IPA software Ingenuity Pathway Analysis (IPA) 01.12 (Qiagen Bioinformatics) to assess canonical pathway analyses. Overrepresentation analysis (ORA) was conducted using WEB-based Gene SeT AnaLysis Toolkit (WebGestalt; www.webgestalt.org^50^). For heatmap representation of selected genes and pathway activation z-scores Morpheus (https://software.broadinstitute.org/morpheus) was used. Cell type expression signature lists were generated from published datasets of mouse RPE/choroid^51^ and retina^52^ single-cell profiling studies. These lists were used for bulk RNAseq deconvolution using an *in-silico* cytometry approach, CIBERSORTx^53^, to estimate cell type representation in the extracted RNA samples of each tissue.

### Data statement

The entirety of the sequencing data are available for download in FASTQ format from NCBI’s Gene Expression Omnibus (to be included on acceptance). Supplemental data files supporting all transcriptomic analyses (Figures 7-12; Supplemental Figures 1-5) are available in this submission. Other data that support the findings of the study are available from the corresponding authors upon request.

### Statistics and Reproducibility

ERG, OKT, and morphometry datasets were statistically analyzed using GraphPad Prism version 10.6.1 (San Diego, CA) and represented as dot plots with underlying bar graphs, line graphs, bar graphs, with mean ± SEM (standard error of the mean) or interleaved box plot & whiskers graphs. For transcriptomic data statistical analyses, the Strand NGS (Version 4.0) built-in analytical pipeline was used, and GraphPad Prism was used for visualization of select gene expression comparisons or pathway/over-representation analyses.

## Results

The Nrf2 KO murine model has been used extensively in preclinical models of ocular disease, particularly to understand the effect of increased oxidative stress susceptibility on retinal/RPE degeneration. Despite important contributions addressing tissue function, morphology, and biochemistry of the retina and RPE in response to combined Nrf2 ablation and stress injury, the interaction of Nrf2 deletion with normal aging has not been comprehensively characterized at the functional or molecular level. The present study was designed to contextualize visual acuity, electroretinographic responses, and retina structure assessments with the transcriptomic profiles of retina and RPE/choroid eyecups of Nrf2 WT and Nrf2 KO mice at 3 mo (young) and 14 mo (adult), across sexes **(Fig. 1)**. The selection of 14 mo as the aged time point in this study was aimed at capturing transcriptional changes in both RPE-choroid and retinal interfaces putatively related to the pathogenesis of early ocular phenotypes in Nrf2 KO mice, described to be apparent in the 11-15 months of age range^32,54^.

**Fig 1.**
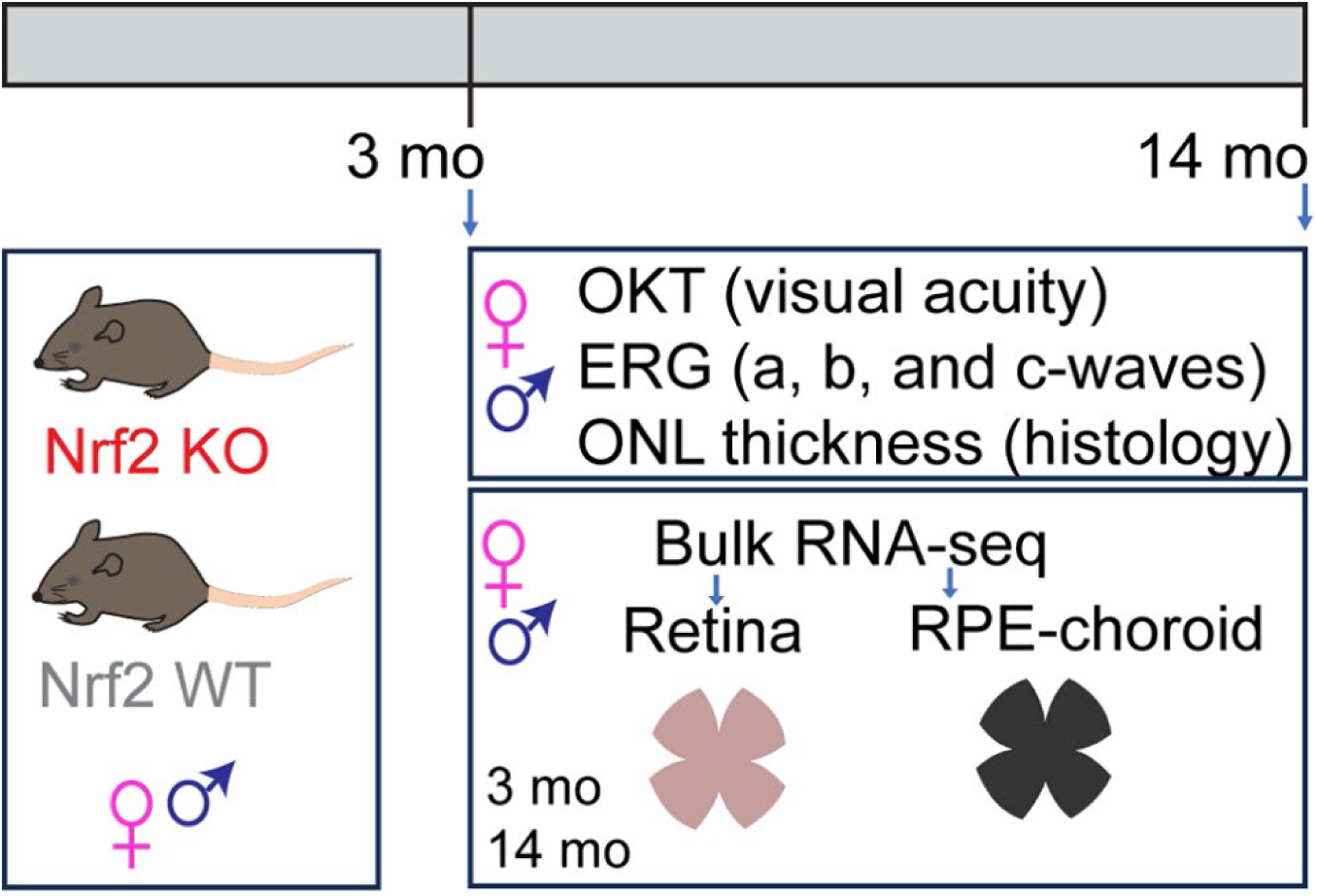
Study design. Female and male Nrf2 null (Nrf2 KO) and WT mice we used at 3 mo and 14 mo for the comparative analysis of visual acuity and electroretinography (ERG) response, and eyes harvested for retina morphometry analyses (ONL thickness measurement) or for transcriptome studies of their retinas and RPE-choroid eyecup tissues by bulk-RNAseq.

### Visual function assessment in young and adult Nrf2 KO and WT mice

Global Nrf2 deletion has been linked to significant exacerbation of visual acuity deficits in the experimental autoimmune encephalomyelitis (EAE) model^30^, but how Nrf2 expression status affects visual function in response to aging has not been reported. To assess visual acuity in response to Nrf2 deletion and age, male and female Nrf2 WT and Nrf2 KO mice were tested at 3 mo and 14 mo for their optokinetic response **(Fig. 2)**. The analysis showed a significant age-related decline in visual acuity for each genotype which was accentuated in the Nrf2 KO group **(Fig. 2a)**, as evidenced by an interaction effect of age and genotype. We found the age-related reduction of visual acuity was maintained when analyzing females and males within Nrf2 WT **(Fig 2b)** and Nrf2 KO **(Fig. 2c)** genotype groups. In the aged Nrf2 WT and Nrf2 KO groups, females had a modest reduction in visual function relative to their male counterparts **(Fig. 2b-c)** but no interaction effects were found between age and sex.

**Fig 2.**
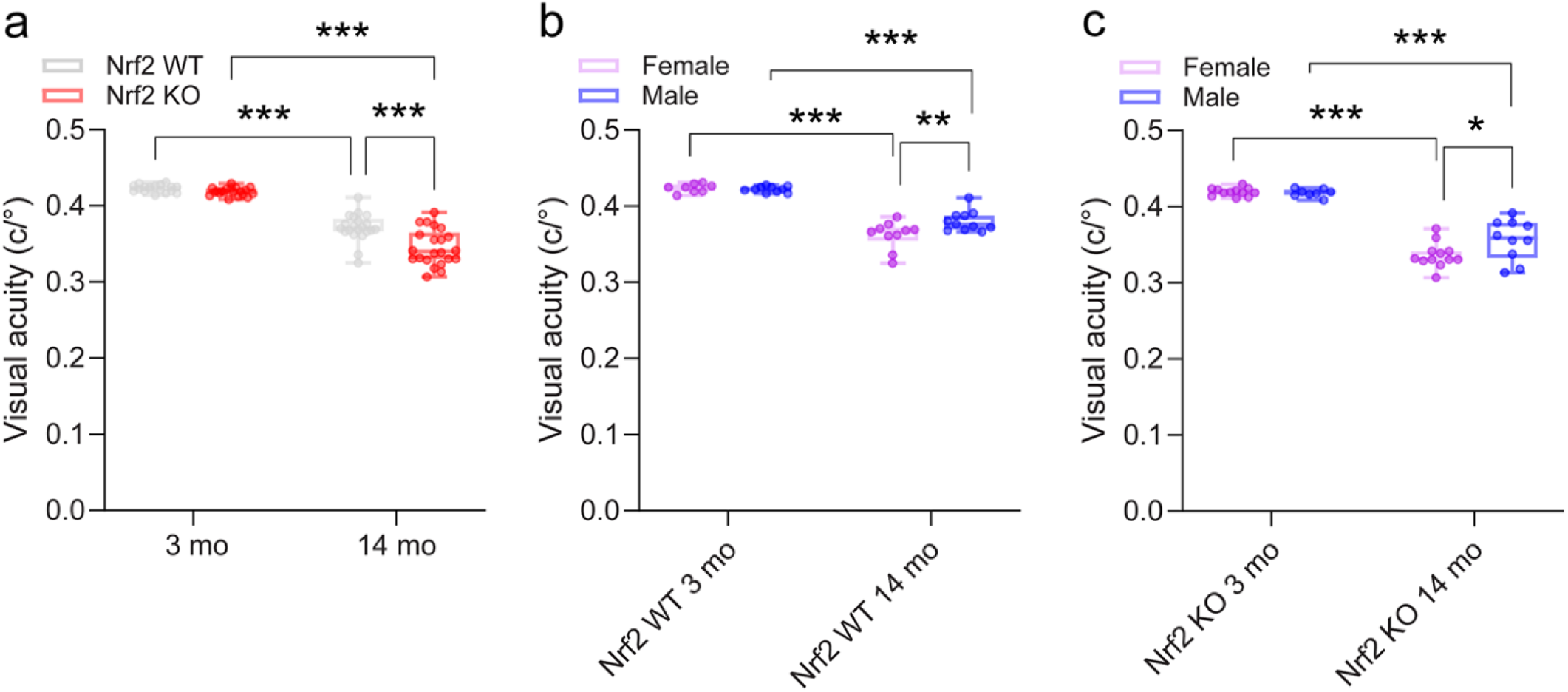
Visual function assessment in young and adult Nrf2 KO and WT mice. Visual acuity was measured in male and female Nrf2 KO and WT mice using OKT optokinetic tracking (OKT) responses at 3 mo and 14 mo. **a:** Interleaved box & whiskers plots depict min to max OKT responses for each genotype and age group. **b-c:** Bar graphs depict averaged OKT responses for each sex and age within each genotype group (b: Nrf2 WT and c: Nrf2 KO). n=18-22/age/genotype in (a); n=8-11/sex/age for the Nrf2 WT group in (b); n=8-12/sex/age for the Nrf2 KO group in (c), *, **, *** p<0.05, p<0.01, and p<0.001, respectively by ordinary 2-way ANOVA with factors age and genotype in (a) and factors age and sex in (b-c), with Tukey’s multiple comparison test.

### Electroretinographic responses of Nrf2 WT and Nrf2 KO mice as a function of age

Electroretinography measures the collective electrical activity of the retina and RPE in response to light stimuli. This measure of retinal function is used both clinically and pre-clinically in models of retinal disease and ocular therapeutics^46,55^. Aging is a biological variable known to affect ERG amplitudes in humans and mice^43,46,56–59^. To assess rod photoreceptor and RPE function, ERG responses evoked by brief, full-field light flashes are recorded under dark-adapted conditions. The relatively rapid a- and b-wave responses, which reflect primarily the activity of photoreceptors and bipolar cells, respectively, are followed by a series of slow potentials generated in response to neural activity by nonneuronal elements of the retina. The c-wave is formed by two components, consisting of hyperpolarization of the RPE basolateral membrane in response to the light-induced decline in subretinal [K^+^], due to photoreceptor activity, and a separate contribution originating from MGC activity^60,61^. As for visual acuity, the literature on ERG responses in the Nrf2 KO mouse model is scarce^32,62^, with one study showing decreased scotopic a- and b-wave responses to a single flash of light in Nrf2 KO compared to Nrf2 WT mice at 12 mo^32^, and no reports on c-wave responses. To address if the visual acuity declines with aging observed in Nrf2 KO mice **(Fig. 2)** are associated with a decrease in electrical responses of the retina and/or RPE, we recorded ERG responses from dark-adapted 3 mo and 14 mo male and female Nrf2 WT and Nrf2 KO mice. As shown in **Fig. 3a-b**, scotopic a- and b-amplitudes were measured in response to light stimuli for all genotypes and ages. Consistent with published reports^43,46,55,57,63^, within each genotype, we observed an age-dependent reduction in the amplitude of a- and b-wave amplitudes. Of interest, a genotype effect was observed at the younger age (3 mo), with Nrf2 KO having significantly lower a- and b-wave amplitude responses compared to their WT counterparts, but this difference in ERG responses was no longer sustained at 14 months of age. Consistent with previous observations, the a-wave (Ta) **(Fig. 3d)** and b-wave (T_b_) **(Fig. 3e)** implicit times at each light stimulus showed no statistical differences between groups^43,64^. Stratifying the analyses by age and sex **(Fig. 4)**, we found that the age-related declines in a- and b-wave amplitudes were maintained within male and female Nrf2 WT **(Fig. 4a-4c)** and Nrf2 KO **(Fig. 4b-4d)** mice but no interaction effects between age and sex were observed.

**Fig 3.**
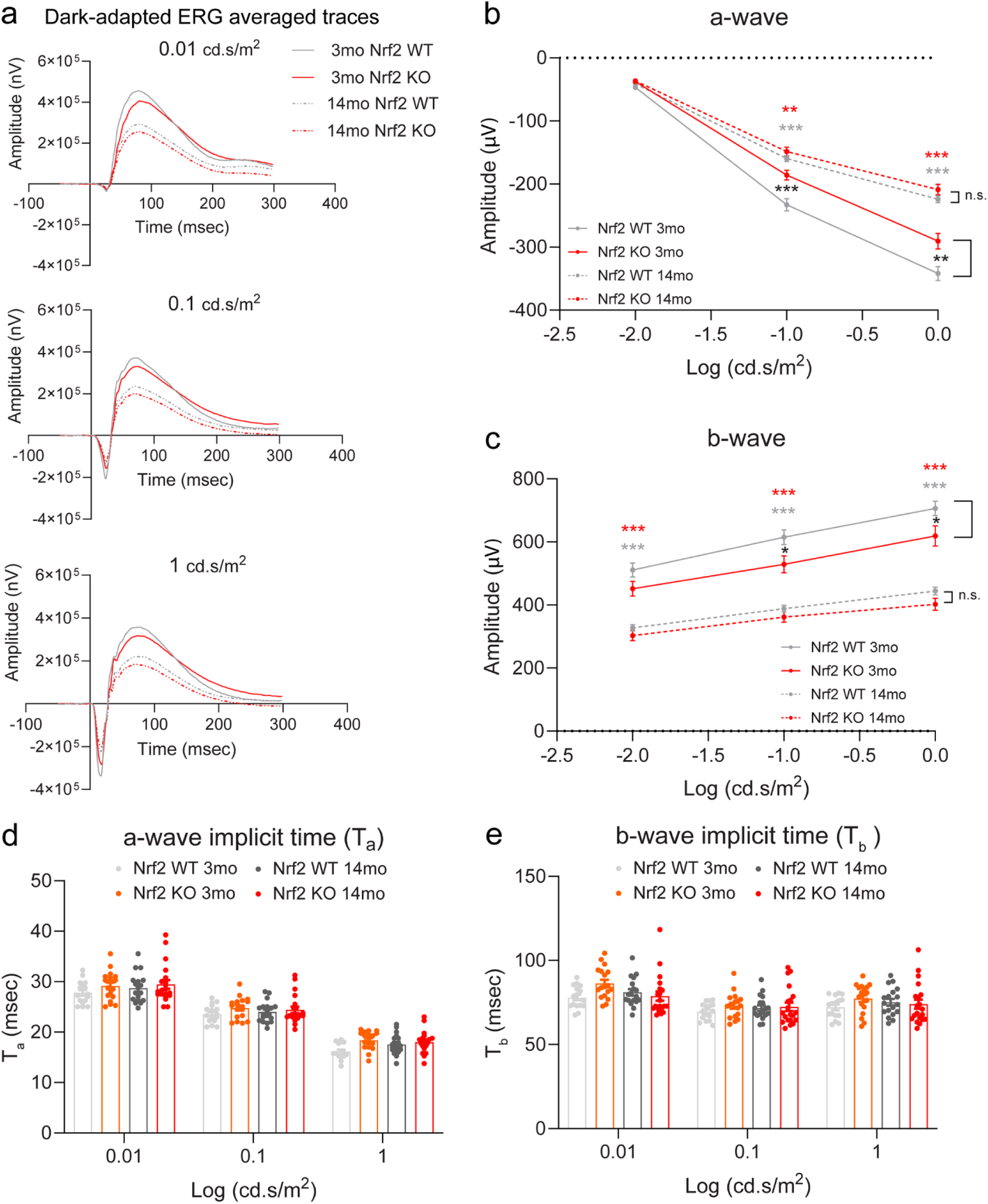
Scotopic a- and b-wave ERG responses of Nrf2 WT and Nrf2 KO mice as a function of age. At 3 mo and 14 mo, Nrf2 WT and Nrf2 KO mice were dark-adapted overnight and subjected to ERG recordings of scotopic a- and b-wave responses to flashlight intensities. a: Representative dark-adapted flash waveforms elicited by increasing luminance flashes (0.01, 0.1, and 1.0 cd.s/m^2^) for each age and genotype group. Line graphs depict the averaged (b) photoreceptor-driven response (a-wave amplitude) and (c) bipolar/Müller cell-driven response (b-wave amplitude) at the different intensity flashes in each group. d-e: Averaged implicit time for (d) a-waves and for (e) b-waves are represented for each intensity flash in each genotype and age group. n=16-20/age/genotype, Mean ± SEM. Red asterisks depict 14 mo Nrf2 KO vs 3 mo Nrf2 KO, gray asterisks depict 14 mo Nrf2 WT vs 3mo Nrf2 WT, black asterisks depict 3mo Nrf2 KO vs 3 mo Nrf2 WT comparisons. *, **, *** p<0.05, p<0.01, and p<0.001, respectively by ordinary 2-way ANOVA with factors age and genotype, with Tukey’s multiple comparison test.

**Fig 4.**
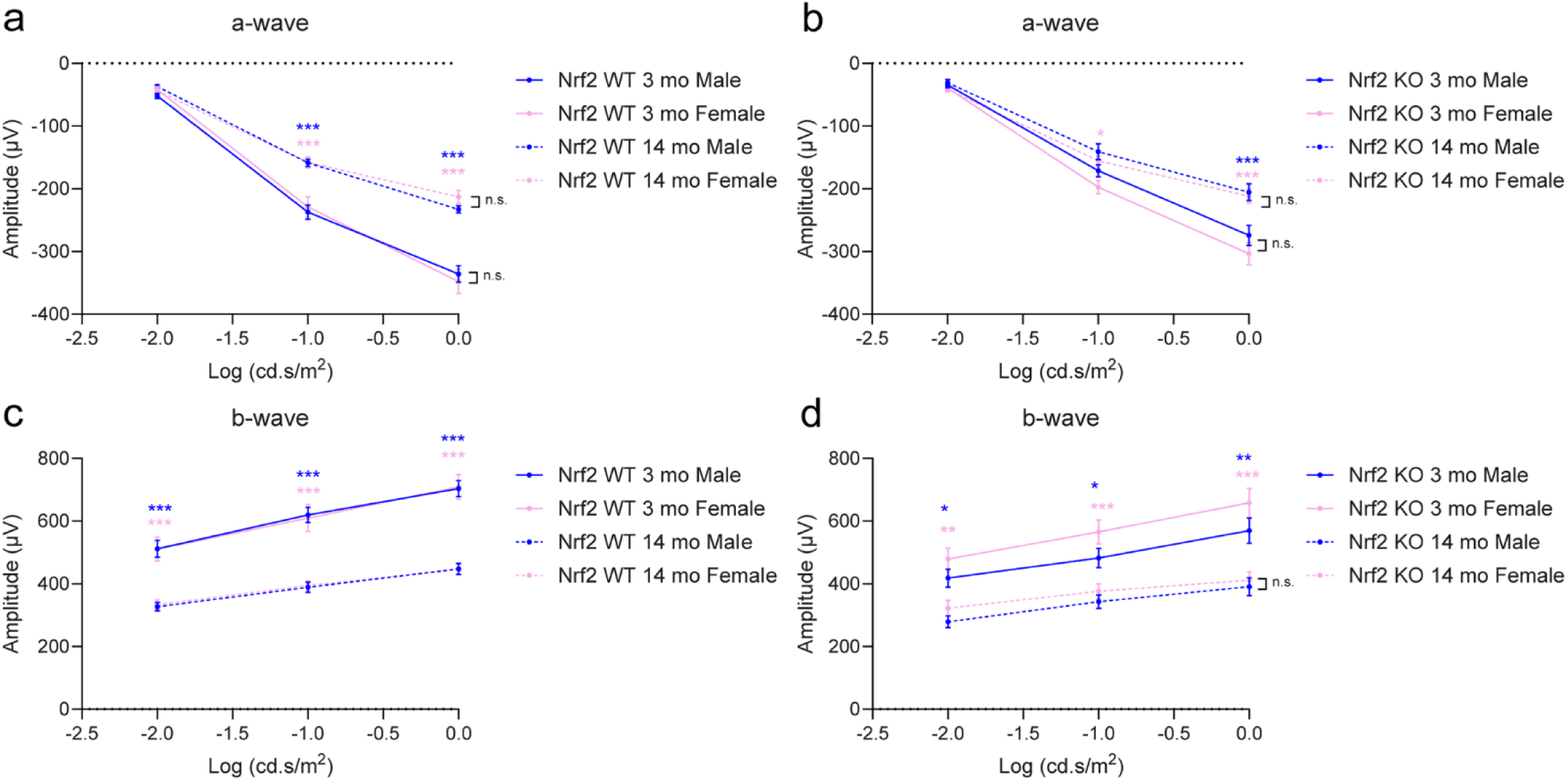
Scotopic a- and b-wave ERG responses by sex and age. The analysis of dark-adapted a- and b-waves from Fig. 3 was stratified by sex and age within each genotype group. **a-b:** Line graphs depict the averaged a-wave amplitude in Nrf2 WT and Nrf2 KO mice with age and sex at the different intensity flashes. **c-d:** Line graphs depict the b-wave amplitude in Nrf2 WT and Nrf2 KO mice with age and sex at the different intensity flashes. n=7-11/sex/age for the Nrf2 WT group and n=8-11/sex/age for the Nrf2 KO group, Mean ± SEM. Blue and pink asterisks depict male and female comparisons, respectively. *, **, *** p<0.05, p<0.01, and p<0.001, respectively by ordinary 2-way ANOVA with factors age and sex, with Tukey’s multiple comparison test.

C-wave analysis revealed that the RPE-evoked electrical signal was reduced with aging and genotype, as visualized by representative waveforms **(Fig 5a)**. Analysis of c-wave maximum response to a single light stimulus revealed a significant amplitude reduction in older mice compared to younger counterparts in each genotype group, but the amplitude reduction at 14 mo was significantly greater in the Nrf2 KO group than in the Nrf2 WT group **(Fig. 5b)**. We previously reported reduced and delayed c-wave response in 24 mo mice relative to 6 mo mice^43^. In the present study, comparing 14 mo vs 3 mo mice, the implicit time for the c-wave response was not delayed with aging or genotype **(Fig. 5c)**. Stratifying the analyses by age and sex **(Fig. 5d-e)**, we found that the age-related reduction in c-wave amplitudes was maintained within male and female Nrf2 WT **(Fig. 5d)** and Nrf2 KO **(Fig. 5e)** mice but no interaction effects between age and sex were observed.

**Fig 5.**
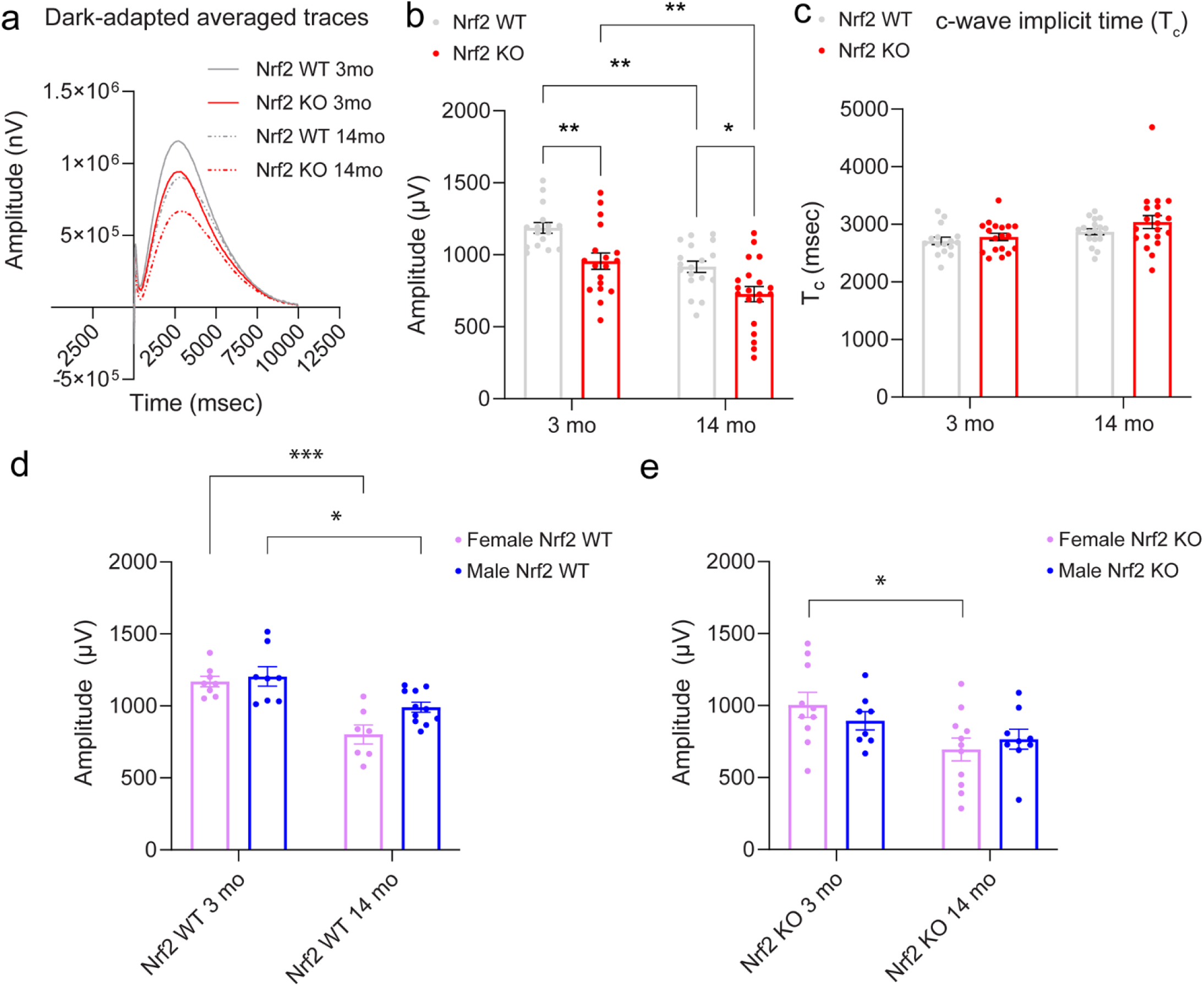
c-wave ERG responses Nrf2 KO and WT mice as a function of age. a: Representative waveforms of RPE-driven c-wave responses to a single light flash (0.9 cd.s/m^2^) from Nrf2 WT and Nrf2 KO mice at 3 mo and 14 mo. **b:** Bar graphs depict the average c-wave amplitudes for each age and genotype group. **c:** Averaged implicit time c-wave is represented for all age and genotype groups. **d-e:** Dark-adapted c-wave responses stratified by sex and age within each genotype group. Bar graphs depict the averaged c-wave amplitude in (d) Nrf2 WT and (e) Nrf2 KO mice with age and sex at the presented intensity flash. b-c) n=16-20/age/genotype, d-e) n=7-11/age/sex in the Nrf2 WT group and n=8-11/age/sex in the Nrf2 KO group. Mean ± SEM. *, **, *** p<0.05, p<0.01, and p<0.001, by (b-c) 2-way ANOVA with age and genotype as variables and (d-e) 2-way ANOVA with age and sex as variables, followed by Tukey multi-comparison test.

### Retina outer nuclear layer thickness analysis of Nrf2 WT and Nrf2 KO mice with age and sex

To assess retina structure in response to age and Nrf2 deletion, eyeballs were harvested from Nrf2 WT and Nrf2 KO mice following ERG recordings and processed for hematoxylin-eosin-stained sagittal tissue sections **(Fig. 6)**. Retina morphometric analysis was performed by measuring ONL thickness at intervals from the ON to the peripheral aspects of the eye. The quantitative analysis showed isolated distance points from the ON (0.6, 1.6, 2.0, and 2.2 mm) in which ONL thickness reduction with age was measured in the Nrf2 WT, Nrf2 KO, or both genotype groups **(Fig. 6i),** but no significant thickness differences were observed between genotype groups. The statistical significance of thickness differences with age **(Fig. 6i)** were lost when stratifying the samples by age and sex **(Fig. 6j)**.

**Fig 6.**
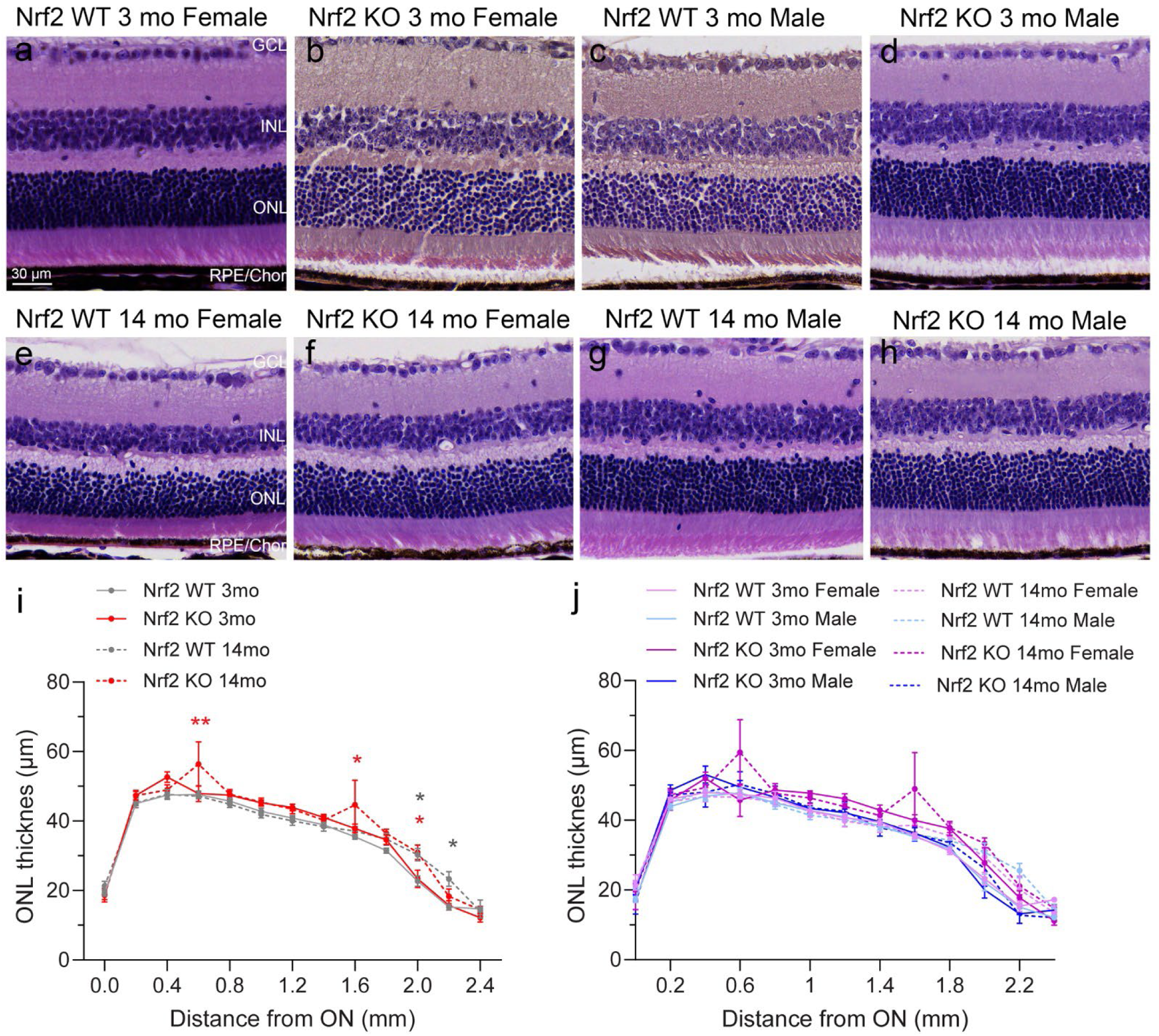
Retina outer nuclear layer thickness analysis of Nrf2 WT and Nrf2 KO mice with age and sex. Following ERG recordings at 3 mo and 14 mo, eyeballs were enucleated from Nrf2 WT and Nrf2 KO mice and processed to generate hematoxylin-eosin-stained sagittal tissue sections. Photoreceptor cell layer thickness, measured as the thickness of the ONL, was assessed from captured scans of the whole tissue sections, at equidistant points from the optic nerve to the periphery of both hemispheres. **a-h:** Representative images of the retina structure from female and male Nrf2 WT and Nrf2 KO mice from both age groups. **i-h:** “Hemi”-spidergraphs were plotted by averaging the ONL thickness values from both eye hemispheres for each group. Analysis is shown with (i) age and genotype and (h) age and sex as variables within each genotype. Data are presented as mean ± SEM, n=12-16/age/genotype in (i) and n=4-8/age/sex in (i). *, ** p<0.05, p<0.01 by 2-way ANOVA with Tukey multiple comparison test.

**Fig 7.**
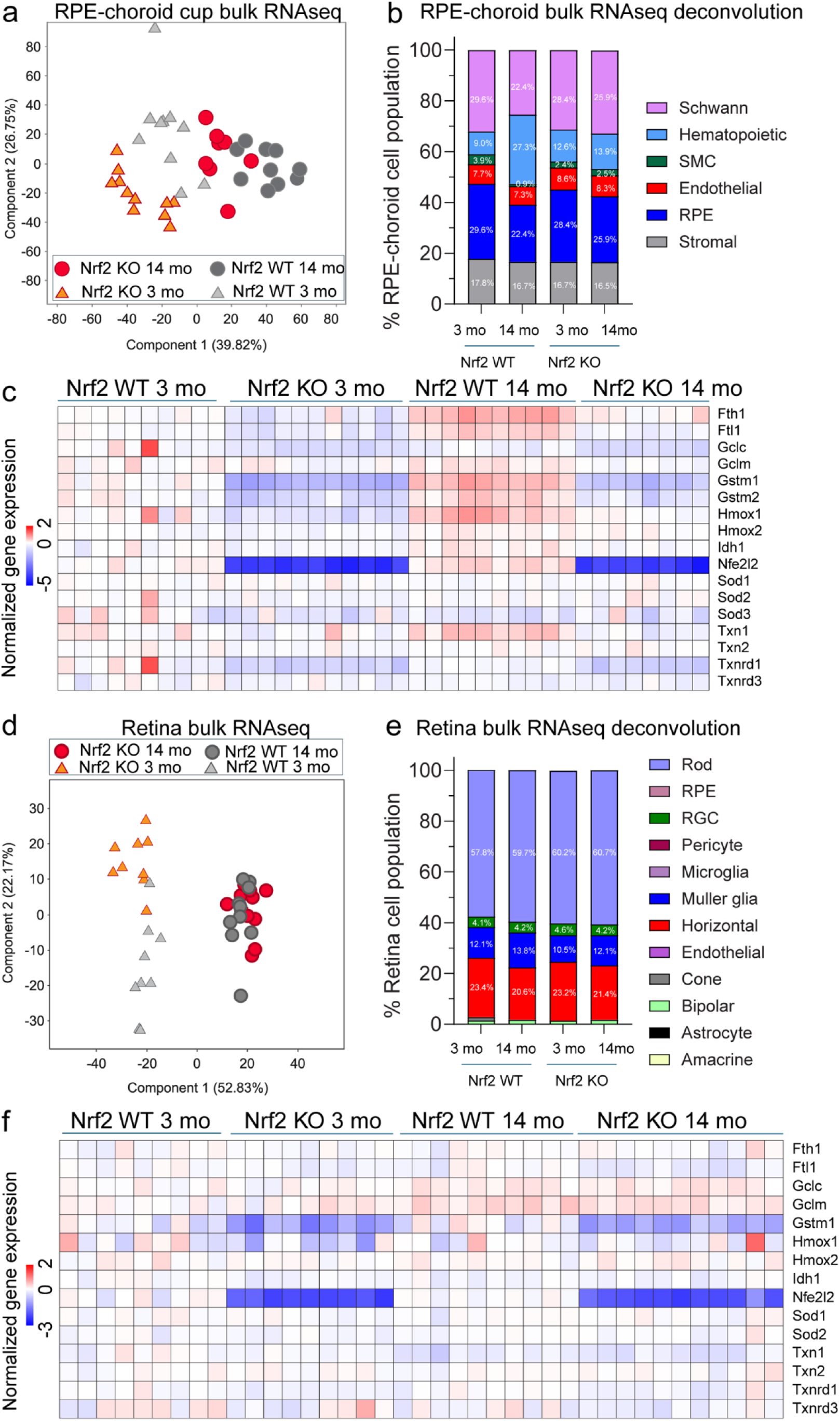
Gene expression profiles of RPE-choroid and retina samples. Following ERG recordings at 3 mo and 14 mo, eyeballs from Nrf2 WT and Nrf2 KO were harvested and dissected for bulk RNA isolation and downstream library construction and stranded RNA sequencing of RPE-choroid and retina tissues. **a:** Principal component analysis (PCA) of expressed genes in RPE-choroid samples showed separation of samples by age and genotype in the first and second components. **b:** Deconvolution of RNAseq data was performed via CIBERSORTx, using scRNAseq mouse eye choroid datasets as reference *(Lehmann et al., JAMA, 2020)*, to estimate cell type representation within the bulk transcriptome of RPE-choroid samples. **c:** Heatmap visualization shows expression of Nrf2 related genes in each RPE-choroid group, including the expected depletion of the Nrf2 gene (*Nfe2l2*) itself in the Nrf2 KO groups while modest alterations in the expression of downstream targets of Nrf2 at both ages, compared to Nrf2 WT counterparts. **d:** Principal component analysis (PCA) of expressed genes in retina samples showed separation of samples by age and genotype in the first and second components. **e:** Deconvolution of RNAseq data was performed via CIBERSORTx, using scRNAseq mouse retina datasets as reference *(Li et al., iScience, 2024)*, to estimate cell type representation within the bulk transcriptome of retina samples. **f:** Heatmap visualization shows expression of Nrf2 related genes in each retina group, including the expected depletion of the Nrf2 gene itself in the Nrf2 KO groups while modest alterations in the expression of downstream targets of Nrf2 at both ages, compared to Nrf2 WT counterparts.

### Gene expression profiles of RPE-choroid and retina samples

To reconcile the functional changes in visual acuity **(Fig. 2)** and scotopic ERGs **(Figs. 3-4-5)** with the apparent absence of gross morphological retinal changes **(Fig. 6)**, we next pursued transcriptomic profiling of the retina and RPE-choroid as a function of age and Nrf2 expression status. To date, how normal aging regulates gene expression of cell types in the retina and RPE-choroid interface to contribute to the pathogenesis of AMD is an outstanding question, for which complimentary bulk and cell-specific level RNA sequencing approaches can provide insights. Following ERG recordings at 3 mo and 14 mo, the contralateral eye to that used for retina ONL thickness measurements **(Fig. 6)** was harvested from each Nrf2 WT and Nrf2 KO mouse and dissected out to obtain a retina and an RPE-choroid cup sample. RNA isolation, downstream library construction, and downstream stranded RNA sequencing of RPE-choroid and retina tissues were performed to interrogate the bulk-tissue level transcriptomic response to Nrf2 deletion and aging in each tissue **(Fig. 7, Supp. data 1)**. RNAseq analysis of expressed genes in RPE-choroid samples, as visualized by Principal Component Analysis (PCA), showed separation of transcriptomes by age and genotype in the first and second components **(Fig. 7a, Supp data 1)**. To estimate cell type representation within the bulk transcriptome of RPE-choroid samples, a deconvolution approach was applied via CIBERSORTx^53^, using scRNAseq mouse eye choroid datasets as reference^51^ **(Fig. 7b, Supp data 1)**. The deconvolution analysis revealed consistent cell representation of RPE (∼23-30%), “Schwann” (∼23-30%), hematopoietic (∼9-27%), stromal (∼17%), endothelial (∼6-9%), and smooth muscle (∼1-4%) cell expression signatures in the bulk RNA extracts across experimental groups **(Fig. 7b)**. Heatmap visualization of normalized expression of Nrf2 (*Nfe2l2*) and its downstream targets showed the expected ablation of Nrf2 transcript in all Nrf2 KO samples compared to Nrf2 WT counterparts regardless of age. Also, as expected in the absence of a combinatorial injury stimulus, only subtle expression changes in downstream target genes were observed. Of interest, in Nrf2 WT RPE-choroid, the directionality of the aging expression changes (Nrf2 WT 14 mo vs Nrf2 WT 3 mo) was consistent with an amplified Nrf2 target gene stress response. This expression pattern was absent from the Nrf2 KO RPE-choroid sample comparison (Nrf2 KO 14 mo vs Nrf2 KO 3 mo) **(Fig. 7c, Supp data 1)**.

RNAseq analysis of expressed genes in retina samples showed segregation of transcriptomes by age and genotype in the first and second components **(Fig. 7d, Supp data 1)**. Of note, the separation of expression profiles by genotype was evident at 3 mo but not at 14 mo. These molecular data were consistent with scotopic ERG a-and b-wave amplitude reductions with genotype at 3 mo but not at 14 mo, indicating a genotype convergent effect with aging on neuroretina cell function and bulk-level gene expression signatures.

The retina is comprised of 11 major cell types, represented by nearly 40 different subpopulations, that collectively capture, process, and transmit electrical signals to the brain^65,66^. The retinal cell population is heavily dominated by photoreceptor neurons (∼80% cell constituency), which is often reflected in mouse^67,68^ and human^6,69^ retina sc-RNAseq studies where rods comprise the main cell cluster recovered. In agreement with these observations, CIBERSORTx deconvolution analysis of our retina expression data across experimental groups, using a mouse single-cell atlas as reference^52^, revealed the greatest transcriptome cell representation was for rod photoreceptors (58-61%), followed by horizontal cell (∼20-23%), MGC (11-14%), RGC (∼4%), and bipolar cell (<2%) expression signatures **(Fig. 7e, Supp data 1)**. Visualization of normalized expression of *Nfe2l2* and its downstream target genes showed the expected ablation of Nrf2 transcript in all retina Nrf2 KO samples compared to Nrf2 WT counterparts regardless of age. Discrete expression changes in downstream target genes of Nrf2 were observed, as visualized by heatmap representation but in contrast to the RPE-choroid **(Fig. 7c)**, the up-regulation of Nrf2 anti-oxidant target genes in response to normal aging was absent in the Nrf2 WT retinas **(Fig. 7f, Supp data 1)**.

### Analytical approach for the study of differential expression profiles of the RPE-choroid and retina in response to aging and Nrf2 deletion

A 2-way ANOVA design with factors age and genotype, with Benjamini-Hochberg multiple testing correction was applied for the study of differential gene expression of RPE-choroid **(Fig 8 a-d)** and retina **(Fig 8e-g)** transcriptomes. The 2-way ANOVA of RPE-choroid samples revealed 1,494 differentially expressed genes (DEGs) with age main effects, 448 DEGs with genotype main effects, and 184 DEGs with age and genotype interaction effects **(Fig. 8a, Supp data 2, Supp. Fig 1)**, as represented by hierarchical clustering **(Fig. 8 b-c-d)**. To identify age-related changes common between or unique to Nrf2 WT and Nrf2 KO RPE-choroid, the 1,494 DEGs with an age main effect were further subjected to one-way ANOVA with SNK pairwise post hoc test. This analysis showed 817 age DEGs in the Nrf2 WT and 1,271 age DEGs in the Nrf2 KO groups with an overlap of 732 DEGs between genotypes. Notably, while only 85 age DEGs were unique to Nrf2 WT RPE-choroid samples, 539 age DEGs were unique to the Nrf2 KO RPE-choroid **(Fig. 8a, Supp data 2)**, which we interpret as indicative of an accelerated aging response in the Nrf2 KO tissue. SNK pairwise post hoc test on the 448 DEGs with a genotype main effect revealed that most of the pair-wise expression changes (151) were captured in the Nrf2 KO vs Nrf2 WT comparisons at both ages. We found 30 genotype DEGs were unique to 3 mo RPE-choroid, and 98 genotype DEGs were unique to 14 mo RPE-choroid transcriptomes **(Fig. 8a, Supp data 2)**.

**Fig 8.**
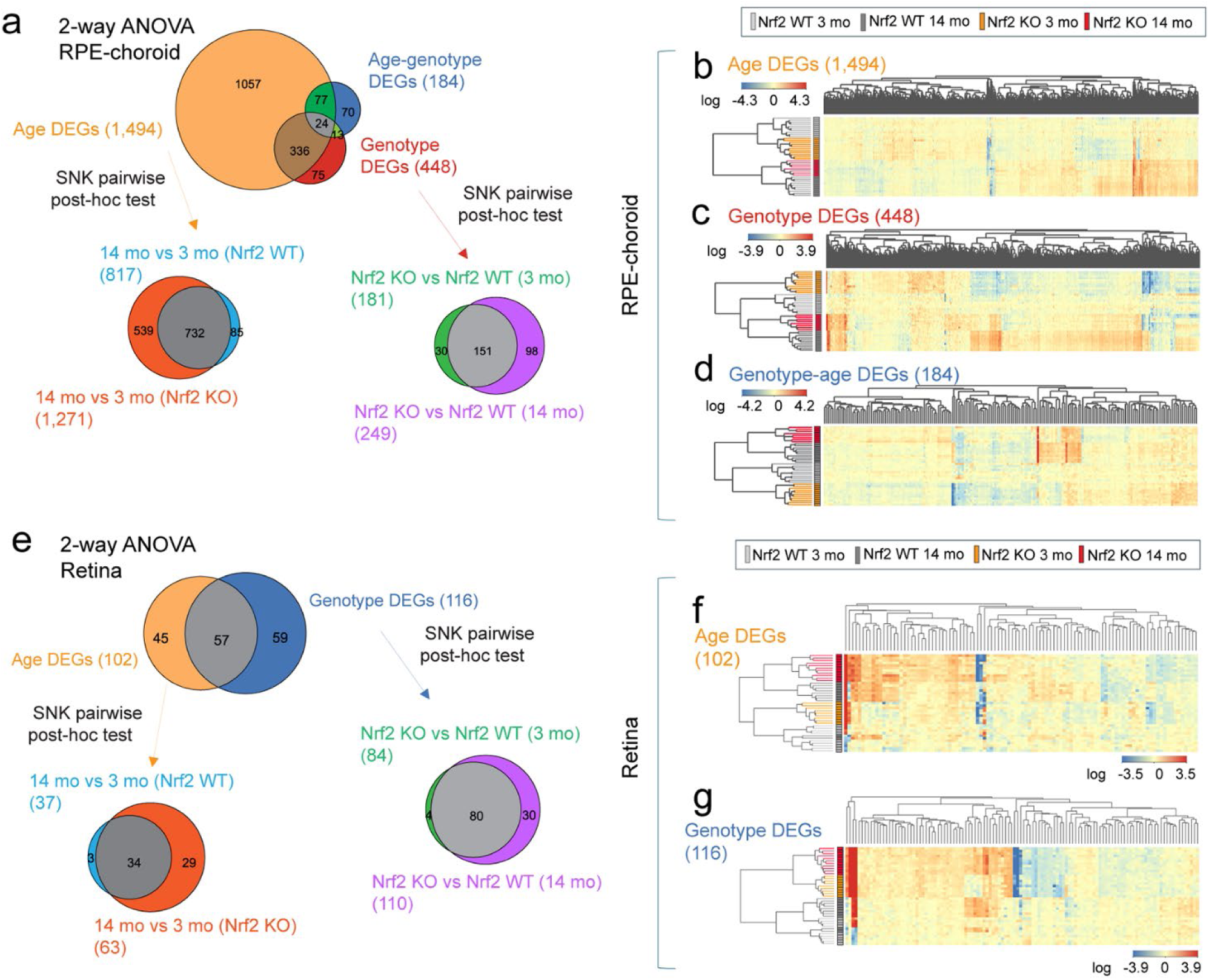
Analytical approach for the study of differential expression profiles of the RPE-choroid and retina in response to aging and Nrf2 deletion. a-d: Age- and genotype-associated DEGs in RPE-choroid samples were identified using 2-way ANOVA with Benjamini-Hochberg multiple testing correction. a: Analysis revealed DEGs with aging (1,494) and genotype (448) main effects, as well as age-genotype interaction effects (184), as visualized by hierarchical clustering in b-c-d. For genes showing a significant age effect (1,494), differences among individual age/genotype groups were then evaluated by 1-way ANOVA with SNK pairwise post-hoc testing and Benjamini-Hochberg correction. SNK of age DEGs revealed age-related changes shared across genotypes (732), unique to Nrf2 KO (539), and unique to Nrf2 WT (85) RPE-choroid samples. For genes showing a significant genotype effect (448), a similar approach was adopted to reveal genotype-related changes shared across ages (151), unique to 3 mo (30), and unique to 14 mo (98) RPE-choroid samples. e-g: Age- and genotype-associated DEGs in retina samples were identified using 2-way ANOVA with Benjamini-Hochberg multiple testing correction. e: Analysis revealed DEGs with aging (102) and genotype (116) main effects, as visualized by hierarchical clustering in f-g. For genes showing a significant age effect (102), differences among individual age/genotype groups were then evaluated by 1-way ANOVA with SNK pairwise post-hoc testing and Benjamini-Hochberg correction. SNK of age DEGs revealed age-related changes shared across genotypes (34), unique to Nrf2 KO (29), and unique to Nrf2 WT (3) retina samples. Analysis of genotype DEGs revealed most genotype-related changes shared across ages (80), unique to 3 mo (4), and unique to 14 mo (30) retina samples. RPE-choroid samples: n=8-11/age/genotype, retina samples: n=8-11/age/genotype.

The 2-way ANOVA of retina samples revealed 102 DEGs with age main effects and 116 DEGs with genotype main effects **(Fig. 8e, Supp data 2)**, as represented by hierarchical clustering **(Fig. 8f-g)**. The interrogation of age DEGs changes within each genotype group by SNK pairwise post hoc test showed that most age DEGs in Nrf2 WT retina (34) were shared with Nrf2 KO retina samples and 29 age DEGs were unique to the Nrf2 KO retina age comparison **(Fig. 8e, Supp data 2)**. SNK pairwise post hoc test on the 116 genotype DEGs revealed that most of the pair-wise expression changes (80) were captured in the Nrf2 KO vs Nrf2 WT retina comparisons at both ages, with 30 genotype DEGs being unique to 14 mo RPE-choroid, and only 4 genotype DEGs unique to 3 mo RPE-choroid transcriptomes **(Fig. 8e, Supp data 2)**.

The data representation in **Fig. 8** reflects a more pronounced differential expression response to aging and Nrf2 deletion in the RPE-choroid **(Fig 8. a-d)** than in the retina **(Fig 8. e-g)**. This observation, along with our ERG findings, is consistent with the widely held notion that during AMD pathogenesis, molecular and functional changes in the choroid-RPE milieu precede degenerative changes in photoreceptor cells^70^.

### Contextualization of the differential gene expression profile of the RPE-choroid in response to aging and Nrf2 deletion

To contextualize the differential expression changes with aging and genotype in the RPE-choroid samples, we conducted pathway analyses **(Fig. 9, Supp data 3)**. Pathway comparison analysis of the 732 age DEGs common to Nrf2 WT and Nrf2 KO RPE-choroid showed preserved predicted activation of immune processes and inhibition of homoeostatic pathways with aging across genotypes within the RPE-choroid milieu **(Fig. 9a, Supp data 3)**. Among the top over-represented pathways were neuroinflammation signaling and phagosome formation, of which normalized expression of associated genes can be observed in **Fig. 9b** and **Fig. 9c**. Consistent with the representation of RPE, endothelial cell, and hematopoietic expression signatures by deconvolution analysis in the bulk transcriptome of RPE-choroid samples **(Fig. 7b)**, the normalized expression of genes collectively associated with innate immune activation of mouse RPE^43^, endothelial^71^, and macrophage^72^ cells under stress, such as *Ccrl2, Ccl5, C3, B2m*, *Syk*, *Clec7a, Cx3cr1*, *Itgam*, and *Tyrobp*, was elevated with age in both Nrf2 WT and Nrf2 KO tissues. Pathways predicted to be inhibited with aging in both genotype groups included collagen biosynthesis/remodeling, extracellular matrix organization, signaling by PDGF, indicative of dysregulation of processes needed to maintain structure of the choroid and retina. Pathway analysis of 539 age DEGs unique to Nrf2 KO RPE-choroid mostly reflected inhibition of fibrotic processes (i.e. pulmonary fibrosis, FAK, wound healing signaling), Notch/Hey1 signaling, and epithelial membrane protein signaling, among others, of which normalized expression of associated genes can be observed in **Fig. 9e-f**. Down regulation of genes coding for Notch receptors (*Notch3*) and ligands (*Dll4, Jag1, Heyl*) with aging has been reported in association with a phenotype of vascular smooth muscle cell drop out from the retina vasculature^73,74^. It is possible that Nrf2 deletion accelerates aging promoting a similar response in the deep choroidal vessels, a phenotype not yet described in the choroidal vascular niche. Pathway comparison analysis of genotype DEGs in RPE-choroid shared across ages showed predicted inhibition of inflammatory and disease responses **(Fig. 9g)**. Normalized expression of genes related to neutrophil degranulation and phagosome formation are shown in **Fig. 9h-i**. Some of these DEGs, up-regulated in the age-comparison within each genotype group (for example, *Clec4d, Clec7a, Tyrobp*) were down-regulated in the Nrf2 KO group compared to WT across ages, thus explaining the differential predicted directionality of pathway activation in the analysis of age DEGs and genotype DEGs. Genotype DEGs unique to 14 mo RPE-choroid were associated with inhibition of neutrophil degranulation, CREB signaling, and molecular mechanisms of cancer **(Fig. 9j-k-l)**. The approach used to analyze differential expressions in RPE-choroid with age and genotype allows us to appreciate that there is a subset of 360 DEGs that overlap across age and genotype main effects **(Fig. 8a)** and that the intersection of 732 age DEGs common to genotype groups and 151 genotype DEGs across ages reveals 132 DEGs shared **(Supp Fig. 2)**. Over-representation analysis of these 132 age and genotype DEGs showed high enrichment ratio for uronic acid metabolism, and processes related to innate and adaptive immune responses **(Supp Fig. 2, Supp data 2)**. Visualization of the expression fold change comparison for most genes involved in these processes further illustrates that pro-inflammatory genes are up-regulated in the 14 mo vs 3 mo comparisons across genotype and down-regulated in the Nrf2 KO vs Nrf2 WT comparisons across ages **(Supp Fig. 2c)**. Of note, Nrf2 has been recently shown to regulate expression of Cd5 antigen-like (CD5L), also known as apoptosis inhibitor expressed by macrophages, with critical functions in inflammation and immunity^75^. In support of this mechanistic axis, the Nrf2 KO RPE-choroid was depleted of *Cd5l*, at both ages studied **(Supp Fig. 2c)**.

**Fig 9.**
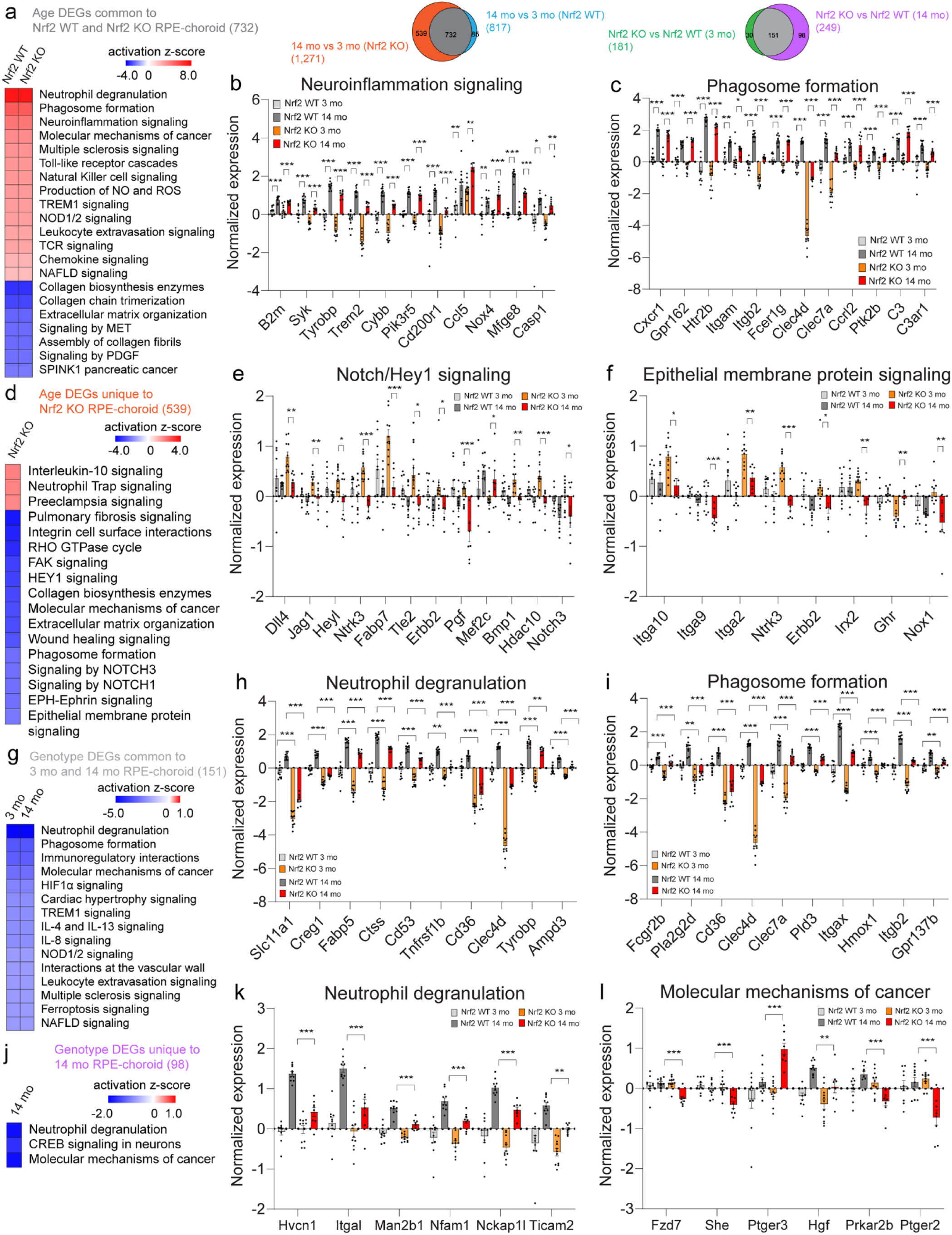
Contextualization of the differential gene expression profile of the RPE-choroid in response to aging and Nrf2 deletion. a-f: Contextualization of expression changes with aging (Fig 8a). a: Pathway analysis of the 732 age DEGs common to Nrf2 WT and Nrf2 KO RPE-choroid showed preserved predicted activation of immune processes and inhibition of homoeostatic pathways with aging across genotypes within the RPE-choroid milieu. b-c: Bars show averaged normalized gene expression for genes in b: neuroinflammation signaling and **b:** phagosome formation for each RPE-choroid group. **d:** Pathway analysis of the 539 age DEGs unique to Nrf2 KO RPE-choroid. **e-f:** Bars show averaged normalized expression for select genes in **e:** Notch/Hey and **f:** epithelial membrane protein signaling. **g-l:** Contextualization of expression changes with genotype (Fig 8a). **g:** Pathway analysis of the 151 genotype DEGs common to 3 mo and 14 mo RPE-choroid showed preserved predicted inhibition of inflammatory signaling processes in Nrf2 KO vs Nrf1 WT across ages. **h-i:** Bars show averaged normalized gene expression for genes in **h:** neutrophil degranulation and **i:** phagosome formation for each RPE-choroid group. **j:** Pathway analysis of the 98 Genotype DEGs unique to14 mo RPE-choroid. **k-l:** Bars show averaged normalized gene expression for genes in **k:** neutrophil degranulation and **l:** molecular mechanisms of cancer for each RPE-choroid group. n=8-11/age/genotype, Mean ± SEM. Age- and genotype associated DEGs were identified by 2-way ANOVA with Benjamini-Hochberg multiple testing correction. For genes showing significant age or genotype effects, differences among individual age/genotype groups were evaluated by 1-way ANOVA with SNK post-hoc testing with Benjamini-Hochberg correction. Adjusted p-values shown in the figure correspond to these post hoc group comparisons (* p<0.05, **p<0.01, and ***p<0.001).

### Contextualization of the differential gene expression profile of the retina in response to aging and Nrf2 deletion

As shown in **Fig 7e**, the differential expression analyses of retina RNA samples yielded a modest number of age and genotype DEGs. Over-representation analysis of the 34 age DEGs shared by Nrf2 WT and Nrf2 KO retinas **(Fig. 10a-b, Supp data 4)** showed enrichment of processes related to odontogenesis, regulation of nervous system, rhythm, and response to metal ion among others, for which normalized expression of associated genes is shown in **Fig. 10g** for all experimental groups. The most comparable process to odontogenesis in the retina is retinogenesis. One of the genes related to this process, *Rspo2*, encodes for R-sponding 2, a critical player in mammalian neuroretina fate differentiation^76^, and was down-regulated in the 14mo retinas compared to 3 mo counterparts in both WT and Nrf2 KO mice. The expression of two genes related to regulation of nervous system, *Myrf* (encodes myelin regulatory factor) and *Trf* (encodes transferrin) increased with age in both Nrf2 WT and Nrf2 KO retinas. Myrf has been described as crucial for retinal development^77^. Trf is an iron transporter not only abundant in human neuroretina and RPE but also significantly up-regulated in the retina of dry AMD patients^78,79^. Over-representation analysis of the 29 age DEGs unique to Nrf2 KO retina showed enrichment of response to toxic substance, renal system development, and symbiotic interaction processes. Normalized expression for some of these unique age DEGs is plotted in **Fig. 10h**.

**Fig 10.**
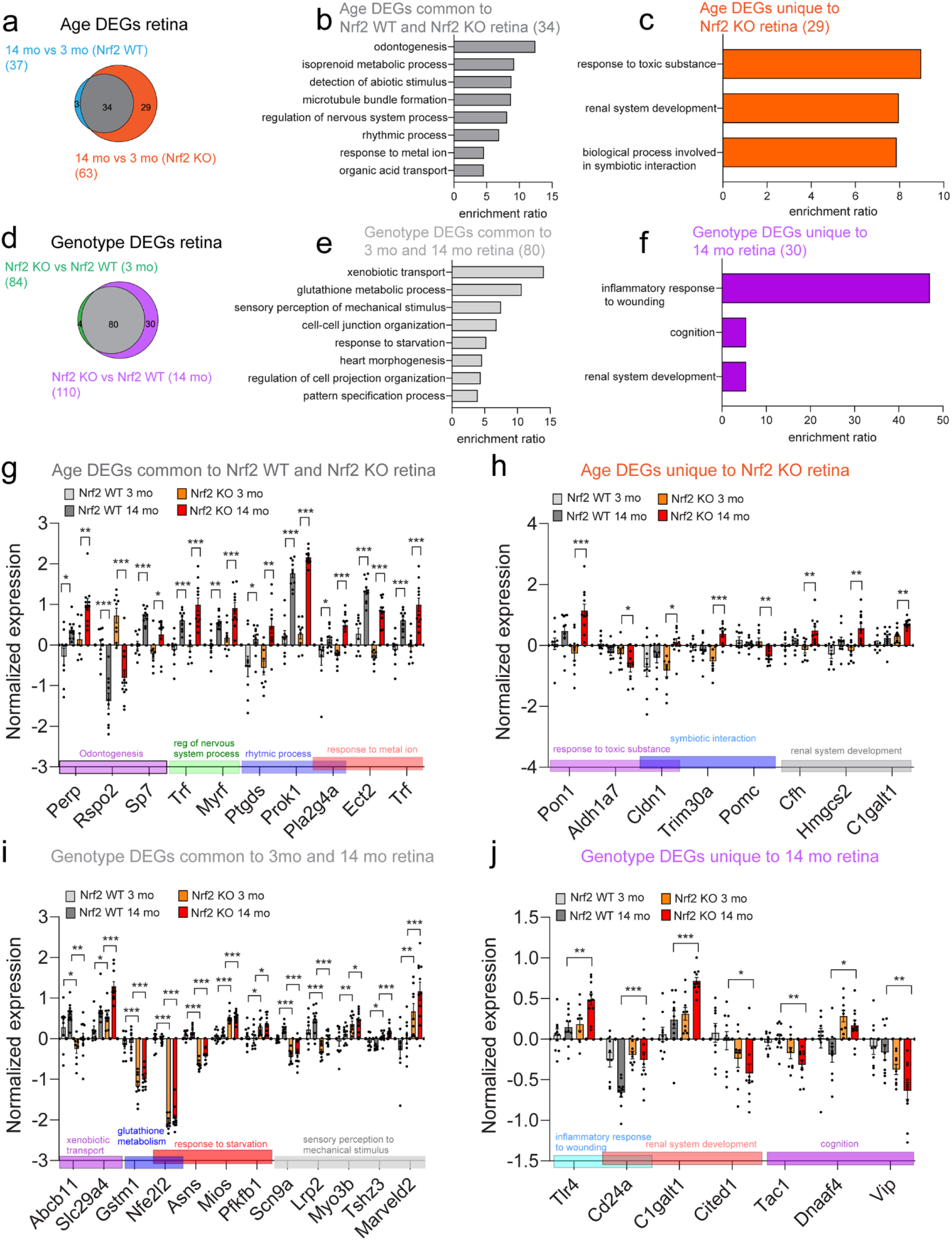
Contextualization of the differential gene expression profile of the retina in response to aging and Nrf2 deletion. a: Contextualization of expression changes with aging (Fig 8e). b-c: Bars depict enrichment ratio for processes identified by overrepresentation analysis (Webgestalt) of **(b)** 31 age DEGs common to Nrf2 WT and Nrf2 KO retina and **(c)** 32 age DEGs unique to Nrf2 KO retina. **d:** Contextualization of expression changes with genotype (Fig 8e). **e-f:** Bars depict enrichment ratio for processes identified by overrepresentation analysis (Webgestalt) of **(e)** 74 genotype DEGs common to 3 mo and 14 mo retina and **(f)** 30 genotype DEGs unique to 14 mo retina. **g-h:** Bars show averaged normalized gene expression for genes in select processes overrepresented in age DEGs **(g)** common and **(h)** unique to age groups. **i-j:** Bars show averaged normalized gene expression for genes in select processes overrepresented in genotype DEGs **(i)** common and **(j)** unique to genotype groups. n=8-11/age/genotype, Mean ± SEM. Age- and genotype associated DEGs were identified by 2-way ANOVA with Benjamini-Hochberg multiple testing correction. For genes showing significant age or genotype effects, differences among individual age/genotype groups were evaluated by 1-way ANOVA with SNK post-hoc testing with Benjamini-Hochberg correction. Adjusted p-values shown in the figure correspond to these post hoc group comparisons (* p<0.05, **p<0.01, and ***p<0.001).

Of interest, the expression of transcripts encoding for complement factor H (*Cfh*), core 1 synthase, glycoprotein-N-acetylgalactosamine 3-beta-galactosyltransferase 1 (*C1galt1*), and HMG-CoA synthase-2 (*Hmgcs2*), under the “renal system development” process, were differentially up-regulated with age only in the Nrf2 KO group **(Fig. 10h)**. Each retinal cell type expresses a specific signature of complement components. Within the murine neuroretina, *Cfh* expression has been assigned mostly to MGC and microglia where its expression decreased with age^66^. Dysregulation of CFH, an important regulatory protein of the alternative pathway of the complement system, has been widely associated with AMD pathogenesis. *Cfh*^-/-^ mice display an ocular phenotype resembling early AMD^80^ and this reported phenotype was rescued by human CFH protein^81^. C1GALT1 has been described for its roles in homeostasis and disease and associated with metastasis and progression of various types of cancer^82^. HMGCS2 is the rate-limiting enzyme controlling ketogenesis and is highly expressed in the RPE where it participates in the production of ketones^83^. Our finding of differential up-regulation of *Hmgcs2* with age in the Nrf2 KO retina but not in the WT counterpart tissue **(Fig. 10h)** suggests a metabolic shift in the aged retina depleted of Nrf2.

The intersection of genotype DEGs from the Nrf2 KO vs Nrf2 WT retina comparisons at 3 mo and 14 mo of age showed most of these DEGs were shared across ages **(Fig. 10d, Supp data 4)**. Over-representation analysis of these 80 genotype DEGs is displayed in **Fig. 10e**, with xenobiotic transport, glutathione metabolism, and sensory perception of mechanical stimulus being the top-3 enriched processes. As expected for the Nrf2 KO mouse, the expression of *Nfe2l2* and its target gene *Gstm1* (encodes glutathione S-transferase) **(Fig. 7f)** in Nrf2 KO retinas was significantly decreased, compared to Nrf2 WT retina samples at 3 mo and 14 mo. Consistent with the idea that lower expression of retinal *Gstm1* in the absence of Nrf2 might be associated with enhanced predisposition to disease, is evidence collected from human post-mortem samples^84^. The mRNA levels of GSTM1 and GSTM5 were significantly reduced in AMD vs age-matched controls in RPE/choroid and neurosensory retina, which was associated with epigenetic repression mechanisms^84^. Apart from glutathione metabolism, *Nfe2l2* was also represented in the response to starvation process in the over-representation analysis.

Only 30 genotype DEGs were unique to the aged Nrf2 KO vs Nrf2 WT retina comparison. The over-representation analysis showed these transcripts were associated with inflammatory response to wounding, renal system development, and cognition **(Fig. 10f, Supp data 4)**. Toll-like receptor (TLR) activation serves as a defense mechanism for the host against infections and tissue damage, initiating a signaling cascade leading to the secretion of various inflammatory cytokines and the activation of immune cells^85,86^. In the mouse retina, a specific member of this family, TLR4, is mainly expressed by microglial cells, MGC, and endothelial cells^87^. In our dataset, retinal *Tlr4* expression was differentially elevated in the Nrf2 KO vs Nrf2 WT retina at 14 mo but not at 3 mo **(Fig. 10j)**, suggesting a combined effect of aging and Nrf2 deletion in the regulation of this pathway. Based on the CIBERSORTx analysis of retina cell-representation **(Fig. 7e)**, in which a signature for MGC but not for endothelial or microglia cell transcriptomes was observed (∼12% across experimental groups), it is possible that the retinal up-regulation of *Tlr4* with aging in our bulk RNAseq is mostly contributed by the MGC compartment. Genotype DEGs unique to the 14 mo retina, under the renal system development process, included *Cd24a*, *C1galt1*, and *Cited1* **(Fig. 10j)**. *C1galt1*, which we identified as an age DEG uniquely in Nrf2 KO retinas **(Fig. 10h)**, was also a genotype DEG, exclusively at the older age. Consistent with a role in oxidative stress, scRNAseq retina studies showed enrichment of *Cd24a* expression in activated MGC clusters following both NMDA treatment and light damage exposure^68^. In our dataset, retinal *Cd24a* was differentially expressed with genotype, exclusively in the aged group **(Fig. 10j)**.

### Comparison of the female and male RPE-choroid transcriptomes in response to aging

We previously published that the transcriptome of the aged retina is sexually divergent^88^ and that the cell-specific translatomic responses of the mouse RPE are age- and sex-divergent^43^. The observation that in the aged Nrf2 WT and Nrf2 KO groups, females had a modest reduction in visual function relative to their male counterparts **(Fig. 2b-c)** further supports the importance of inclusion of sex as a variable in our transcriptomic studies. After assessing the transcriptomic phenotype of the RPE-choroid in response to aging and Nrf2 deletion, we interrogated sex divergences in the RPE-choroid age-related responses within each genotype group. PCA of trascriptomes of Nrf2 KO RPE-choroid tissues showed segregation of expression profiles by age and sex in the first and second components **(Fig. 11a, Supp data 5)**. Two-way ANOVA revealed 260 DEGs with age main effects, 14 DEGs with sex main effects (only 4 of these were autosomal), and no interaction between age and sex effects **(Fig. 11b, Supp data 5)**, as represented by hierarchical clustering **(Fig. 11c-d)**. Of note, the 14 sex DEGs completely overlapped at both ages analyzed **(Fig. 11f, Supp data 5)**. Subjecting the set of age DEGs to one-way ANOVA and SNK pairwise post hoc test showed 154 age DEGs in the female and 118 age DEGs in the male Nrf2 KO RPE-choroid samples with an overlap of ∼49% of these age DEGs across sexes (89) **(Fig. 11e, Supp data 5)**. ∼43% of age DEGs (65) were unique to female and ∼16% (29) were unique to male Nrf2 KO RPE-choroid tissues, suggesting sex-specific signatures with aging. Over-representation analyses of these sets of age DEGs are depicted in **Fig.11 g-h-I (Supp data 5)**. Enriched biological processes in the analysis of age DEGs common to female and male Nrf2 KO RPE choroid were consistent with inflammation, such as acute inflammatory response, regulation of T cell activation, and cytokine-mediated signaling. The normalized expression of select genes associated with these mechanisms is depicted for male and female Nrf2 KO RPE-choroid samples in **Fig. 11j (Supp data 5)**. NPY is a peptide in postganglionic nerves innervating the choroid, relevant for neovascularization, inflammation, and vasoconstriction which has been shown to be-upregulated in diabetic mouse choroids^89^. ORM1 is a plasma glycoprotein associated with inflammation, thought to modulate the permeability of microvessel walls and the blood-brain barrier, it is reported to be increased in the tear film of Alzheimer’s disease patients^90^. Early up-regulation of *Saa3* (encoding serum amyloid A) is a key step in priming inflammation, initiated by macrophages and fibroblasts^91^. *MHC class Ib* (*H2-M2*) encodes a conserved, non-classical major histocompatibility complex (MHC) class I cell surface glycoprotein^92^. The elevated expression of the acute inflammatory response-related genes *Npy*, *Orm1*, *Saa3*, *and H2-M2* with aging in the Nrf2 KO RPE-choroid across sexes, is consistent with dysregulated local immune responses in the absence of Nrf2 signaling. Consistent with the detection of immune compartment cell clusters in the mouse choroid by scRNAseq, such as T cells and macrophages^93^, our bulk RNAseq analysis revealed expression of immune cell signature genes, such as *Cd3e* and *Cd4* (T cells), *Ccl2* and *Mmp12* (macrophages), which increased with aging in female and male Nrf2 KO samples **(Fig. 11j, Supp data 5)**. Over-representation analysis of age DEGs specific for female **(Fig. 11h)** and male **(Fig. 11i)** Nrf2 KO tissues revealed sex-specific changes in immune/inflammatory processes. For example, *Ptgdr* and *Cxcl1* expressions were differentially up-regulated and *Spon2* was down-regulated with age specifically in female Nrf2 KO RPE-choroid. In male but not female Nrf2 KO RPE-choroid, the expressions of *Il1rn*, *Cdo1*, and *Cd14* were differentially up-regulated and *Sigirr* was down-regulated in the older age group **(Fig. 11k, Supp data 5)**.

**Fig 11.**
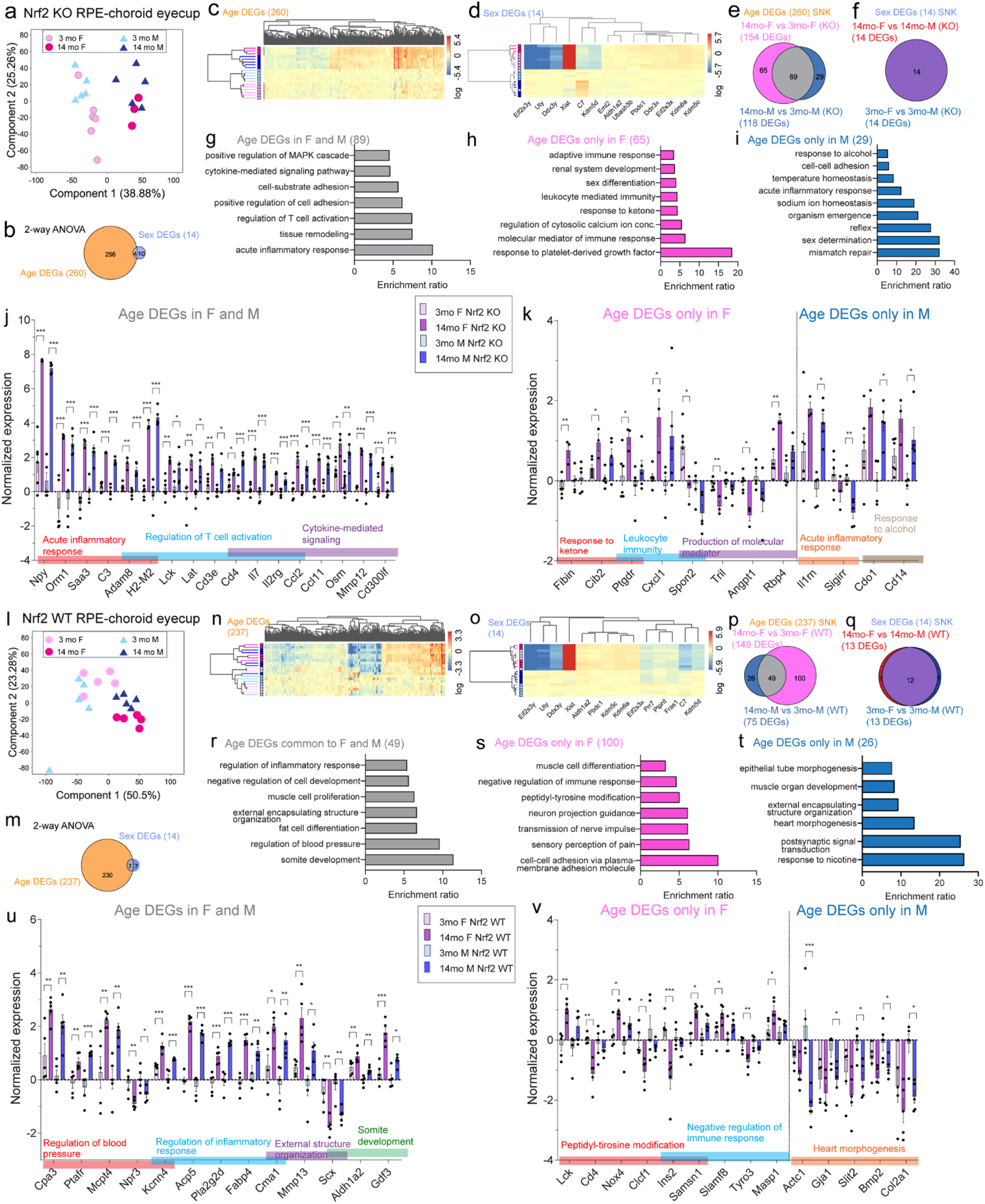
Comparison of the female and male RPE-choroid transcriptomes in response to aging. Age and sex were used as variables to separately analyze the transcriptomic profiles of the Nrf2 WT and Nrf2 KO RPE-choroid groups. **a-k:** PCA of expressed genes showed **(a)** separation of Nrf2 KO samples by age and sex in the first and second components. **b:** 2-way ANOVA with factors age and sex on the RNAseq data revealed DEGs with aging (260) and sex (14) main effects, as visualized by hierarchical clustering in **c**-**d**. **e-f:** SNK pairwise post-hoc test revealed ∼60% age DEGs in Nrf2 KO samples are common across sexes (89) with specific expression signatures in female (65) and male (29) samples, as well as all sex DEGs are common across ages (14). **g-i:** Bar graphs illustrate top-enriched processes identified by over-representation analysis (Webgestalt) of age DEGs common and unique to each sex. **j:** Normalized gene expression for select age DEGs shared by male and female Nrf2 KO RPE-choroid samples in acute inflammatory response, regulation of T cell activation, and cytokine mediated signaling. **k:** Normalized gene expression for select age DEGs unique to female Nrf2 KO and male Nrf2 KO samples from processes from (h) and (i). **l-v:** PCA of expressed genes showed **(l)** separation of Nrf2 WT RPE-choroid transcriptomes by age and sex in the first and second components. **m:** 2-way ANOVA with factors age and sex on the RNAseq data revealed DEGs with aging (237) and sex (14) main effects, as visualized by hierarchical clustering in **n**-**o**. **p-q:** SNK pairwise post-hoc test revealed age DEGs in Nrf2 WT samples common across sexes (49) with specific expression signatures in female (100) and male (26) samples. Most sex DEGs are common across ages (13). **r-t:** Bar graphs illustrate top-enriched processes identified by over-representation analysis of age DEGs in Nrf2 WT samples common and unique to each sex. **u:** Normalized gene expression for select age DEGs shared by male and female Nrf2 WT samples in regulation of blood pressure, inflammatory response, and structure organization/development. **v:** Normalized gene expression for select age DEGs unique to female Nrf2 KO and male Nrf2 KO samples from processes from (s) and (t). n=3-5/age/sex for Nrf2 KO and n=5/age/sex for Nrf2 WT RPE-choroid groups. Mean ± SEM. Age-associated DEGs within each genotype were identified by 2-way ANOVA with Benjamini-Hochberg multiple testing correction. For genes showing significant age effects, differences among individual age/sex groups were evaluated by 1-way ANOVA with SNK post-hoc testing with Benjamini-Hochberg correction. Adjusted p-values shown in the figure correspond to these post hoc group comparisons (* p<0.05, **p<0.01, and ***p<0.001).

**Fig 12.**
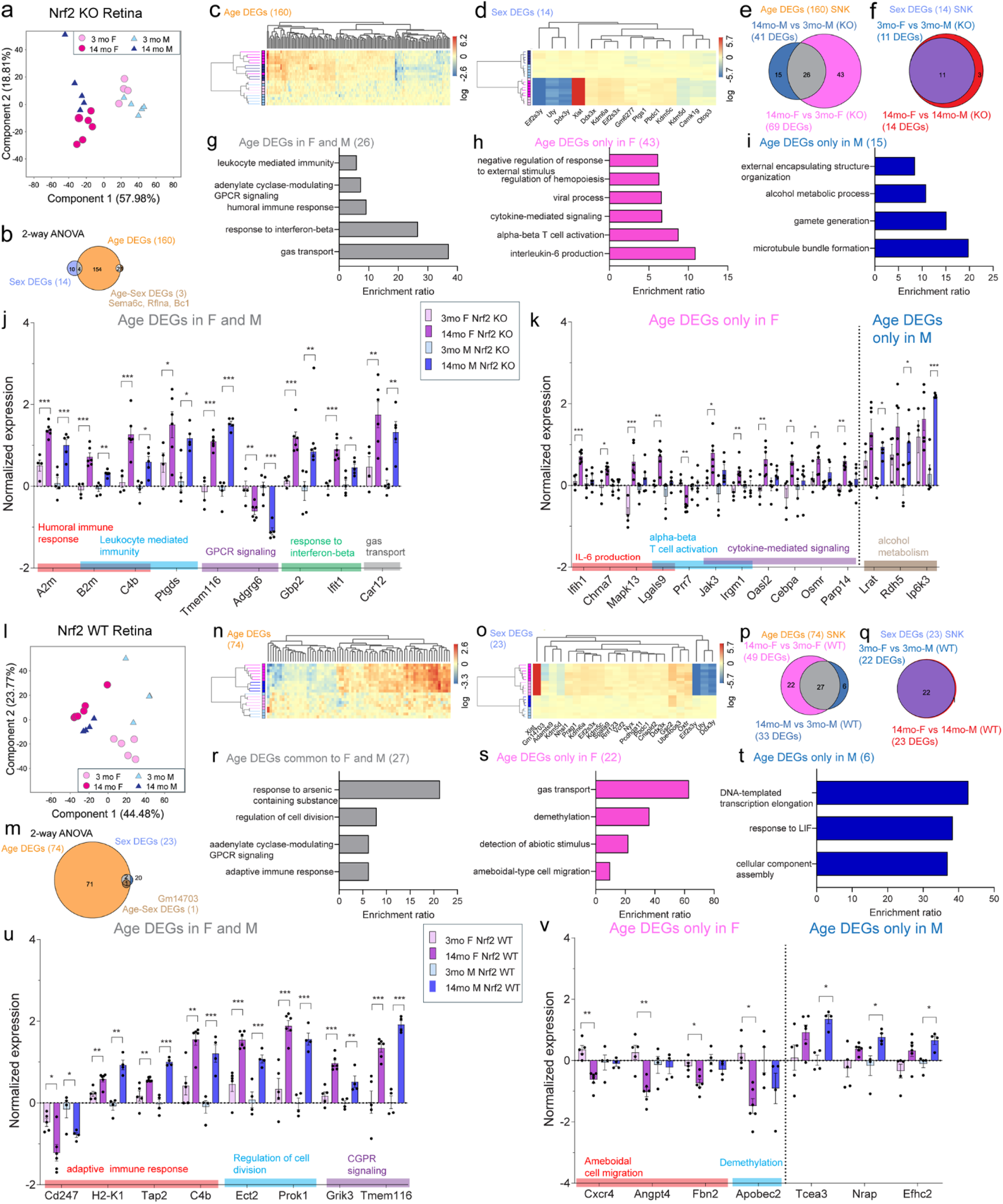
Comparison of the female and male retina transcriptomes in response to aging. Age and sex were used as variables to separately analyze the transcriptomic profiles of the Nrf2 WT and Nrf2 KO retina groups. **a-k:** PCA of expressed genes showed **(a)** separation of Nrf2 KO retina transcriptomes by age and sex in the first and second components. **b:** 2-way ANOVA with factors age and sex on the RNAseq data revealed DEGs with aging (160) and sex (14) main effects, as visualized by hierarchical clustering in **c**-**d**. **e-f:** SNK pairwise post-hoc test revealed a set of age DEGs in Nrf2 KO samples are common across sexes (26) with specific expression signatures in female (43) and male (15) samples, as well as most sex DEGs are common across ages (11). **g-i:** Bar graphs illustrate top-enriched processes identified by over-representation analysis (Webgestalt) of age DEGs common and unique to each sex in Nrf2 KO retina. **j:** Normalized gene expression for select age DEGs shared by male and female Nrf2 KO samples in humoral immune response, leukocyte mediated immunity, response to interferon-β, GPCR signaling, and gas transport. **k:** Normalized gene expression for select age DEGs unique to female Nrf2 KO and male Nrf2 KO samples from processes from (h) and (i). **l-v:** PCA of expressed genes showed **(l)** separation of Nrf2 WT retina transcriptomes by age and sex in the first and second components. **m:** 2-way ANOVA with factors age and sex on the RNAseq data revealed DEGs with aging (74) and sex (23) main effects, as visualized by hierarchical clustering in **n**-**o**. **p-q:** SNK pairwise post-hoc test revealed age DEGs in Nrf2 WT samples common across sexes (27) with specific expression signatures in female (22) and male (6) Nrf2 WT retina samples. Most sex DEGs are common across ages (22). **r-t:** Bar graphs illustrate top-enriched processes identified by over-representation analysis of age DEGs in Nrf2 WT samples common and unique to each sex. **u:** Normalized gene expression for select age DEGs shared by male and female Nrf2 WT retina samples in adaptive immune response, regulation of cell division, and CGPR signaling. **v:** Normalized gene expression for select age DEGs unique to female male Nrf2 WT samples from processes from (s) and (t). n=4-6/age/sex for each Nrf2 KO and Nrf2 WT retina groups. Mean ± SEM. Age-associated DEGs within each genotype were identified by 2-way ANOVA with Benjamini-Hochberg multiple testing correction. For genes showing significant age effects, differences among individual age/sex groups were evaluated by 1-way ANOVA with SNK post-hoc testing with Benjamini-Hochberg correction. Adjusted p-values shown in the figure correspond to these post hoc group comparisons (* p<0.05, **p<0.01, and ***p<0.001).

PCA of Nrf2 WT RPE-choroid transcriptomes showed separation of samples by age and sex in the first and second components **(Fig. 11l, Supp data 6)**. Two-way ANOVA revealed 237 DEGs with age main effects, 14 DEGs with sex main effects (only 4 of these were autosomal), and no interaction between age and sex effects **(Fig. 11m, Supp data 6)**, as represented by hierarchical clustering **(Fig. 11n-o)**. Subjecting the set of age DEGs to one-way ANOVA and SNK pairwise post hoc test showed 149 age DEGs in the female and 75 age DEGs in the male Nrf2 WT RPE-choroid samples with an overlap of 28% of these age DEGs across sexes (49) **(Fig. 11p, Supp data 6)**. We found that ∼57% of these age DEGs (100) were unique to female and ∼15% (26) were unique to male Nrf2 WT RPE-choroid tissues, suggesting a bias for sex-specific signatures with aging in female tissue. Over-representation analyses of these sets of age DEGs are depicted in **Fig.11 r-s-t (Supp data 6)** and normalized expressions of select genes comprised in these biological processes are depicted for female and male Nrf2 WT RPE-choroid at both ages **(Fig. 11 u-v, Supp data 6)**. Related to immunity, a subset of age DEGs (including *Kcnn*, *Acp5*, *Pla2g2*, *Fabp4*, and *Cma1*) **(Fig. 11u, Supp data 6)**, associated with regulation of immune response, was differentially up-regulated with aging in both female and male Nrf2 WT RPE-choroid tissues. Specific to the female sex, was the down-regulation of *Ins2* and *Tyro3*, and up-regulation of *Samsn1*, *Slamf8*, and *Masp1*as part of the negative regulation of immune response process **(Fig. 11v, Supp data 6)**. Over-representation analysis of age DEGs specific to male did not enrich for immune response related processes, but for tissue/organ development, such as heart morphogenesis (see **Fig 11. v** for associated gene normalized expressions), as well as response to nicotine and postsynaptic signal transduction. An observation from the analysis of sex differences that was shared between Nrf2 KO and Nrf2 WT RPE-choroid was differentially higher expression of the complement pathway gene *C7* in male versus female and of *Aldh1l1a2* (encodes aldehyde dehydrogenase 1 family member A2) and *Pbdc1* in female versus male samples (**Fig 11d-11o)**. To determine the extent of overlap of age-related changes in Nrf2 WT and Nrf2 KO female and male RPE-choroid tissues, we intersected the lists of age DEGs in Nrf2 WT and Nrf2 KO female **(Supp Fig. 3a)** and in Nrf2 WT and Nrf2 KO male **(Supp Fig. 3b)** samples (from analyses **in Fig 11e-11p**). Venn representation intersecting the 149 age DEGs in Nrf2 WT and 154 age DEGs in Nrf2 KO female RPE-choroid samples showed only 17 DEGs overlapped between genotypes while 132 and 137 age DEGs were specific for Nrf2 WT and Nrf2 KO genotypes, respectively **(Supp Fig 3a, Supp data 9)**. Similarly, intersection of 75 age DEGs in Nrf2 WT and 118 age DEGs in Nrf2 KO male RPE-choroid showed only 7 DEGs shared across genotypes with 68 and 111 age DEGs being specific for Nrf2 WT and Nrf2 KO genotypes, respectively **(Supp Fig 3b, Supp data 9)**. These comparisons further support that differential expression profiles in response to aging in each sex are influenced by Nrf2 expression status.

### Comparison of the female and male retina transcriptomes in response to aging

Following the same rationale as above for RPE-choroid, after assessing the transcriptomic phenotype of the retina in response to aging and Nrf2 deletion, we interrogated sex divergences in the bulk retina age-related responses within each genotype group. PCA of Nrf2 KO retina transcriptomes revealed segregation of expression profiles by age and sex in the first and second components **(Fig. 12a, Supp data 7)**. Two-way ANOVA revealed 160 DEGs with age main effects, 14 DEGs with sex main effects (only 4 of which were autosomal), and no interaction between age and sex effects **(Fig. 12b, Supp data 7)**, as represented by hierarchical clustering **(Fig. 12c-d)**. Out of the14 sex DEGs 11 overlapped at both ages analyzed **(Fig. 12f, Supp data 7)**. Subjecting the set of age DEGs to one-way ANOVA and SNK pairwise post hoc test showed 69 age DEGs in the female and 41 age DEGs in the male Nrf2 KO RPE-retina samples with an overlap of 31% of these age DEGs across sexes (26) **(Fig. 12e, Supp data 7)**. 51% of age DEGs (43) were unique to female and ∼18% (15) were unique to male Nrf2 KO retina tissues, suggesting sex-specific signatures with aging skewed toward female samples. Over-representation analyses of these sets of age DEGs are depicted in **Fig.12 g-h-I (Supp data 7)**. Most enriched biological processes in the analysis of age DEGs common to female and male Nrf2 KO retina transcriptomes were consistent with inflammation, such as humoral immune response, leukocyte mediated immunity, GPCR signaling, and response to IFN-β, of which normalized expressions for associated genes are depicted in **Fig. 12 j (Supp data 7)**. One of the age-related expression changes in these mechanisms is *A2m* [encodes α2-Macroglobulin (A2M)]. Among its range of functions, the plasma protease inhibitor A2M enhances proliferation and migration of immune and cancer cells (reviewed in ^94^). We and others reported up-regulation of *A2m* in retinal MGC in response to optic nerve crush^39^ and after *Aire* deletion^67^, and in a human immortalized MGC cell line (MIO1-M1) following IL-6 treatment^95^. Consistent with a role in MHC I antigen processing and presentation, components of this pathway have been identified in the neuroretina with aging^96^. We previously identified differential enrichment of the MHC I molecule β2-microglobulin (*B2m*) specifically in MGC^39^, and mouse retina scRNAseq studies have shown its enrichment in the activated MGC subcluster in response to light damage and NMDA treatments^68^. Among all retinal cell types, the MGC compartment, which is represented in our bulk RNAseq deconvolution analysis **(Fig. 7e)**, has been described as the main source of expression of the classical complement pathway component gene *C4b*^66^. Our observation of up-regulation of *A2m*, *B2m*, and *C4b* is consistent with a pro-inflammatory response in both male and female Nrf2 KO retinas with aging **(Fig. 12j)**. Normalized expression for select age DEGs unique to female or male Nrf2 KO retinas, comprised under the biological processes in **Fig. 12h-i** are shown in **Fig. 12k (Supp data 7)**. Unique to female Nrf2 KO tissue was the age-triggered elevation of a set of genes related to IL-6 production, α-β T cell activation, and cytokine-mediated signaling whereas unique to male was the up-regulation of alcohol-metabolism genes with age **(Fig. 12k, Supp data 7)** among others.

PCA of Nrf2 WT retina expression profiles showed segregation of samples by age and sex in the first and second components, with the separation by sex being greater at the 3 mo than 14 mo age **(Fig. 12l, Supp data 8)**. Two-way ANOVA revealed 74 DEGs with age main effects, 23 DEGs with sex main effects (10 of these were autosomal), and no interaction between age and sex effects **(Fig. 12m, Supp data 8)**, as displayed by hierarchical clustering **(Fig. 12n-o)**. Subjecting the set of age DEGs to one-way ANOVA and SNK pairwise post hoc test showed 49 age DEGs in the female and 33 age DEGs in the male Nrf2 WT retinas with 49% of these age DEGs shared across sexes (27) **(Fig. 12p, Supp data 8)**. We found that 40% of these age DEGs (22) were unique to female and ∼11% (6) were unique to male Nrf2 WT retinas. Over-representation analyses of these sets of age DEGs **(Fig.12 r-s-t, Supp data 6)** and normalized expression of select genes comprised in these biological processes are depicted for female and male Nrf2 WT retinas at both ages **(Fig. 12 u-v, Supp data 8)**.

Age DEGs under the adaptive immune response category, sharing directionality of change in male and female Nrf2 WT retina included *Cd247, C4b, H2-K1*, and *Tap2* **(Fig. 12u, Supp data 8)**. Sex specific signatures in the WT retina included decreased *Cxcr4, Angpt4*, and *Fbn2*, captured under “ameboidal cell migration” but collectively related to leukocyte trafficking, modulation of hematopoietic cell proliferation, and choroidal neovascularization. CXCR4 (and its ligand SDF-1) expression has been localized in normal aged human retina and choroid^97^. The angiopoietin ligands (Angpt1-4) signal through the receptor tyrosine kinase Tie1 and Tie2 and Angpt4 was reported to be necessary for venous remodeling and SMC maturation in the mouse retina^98^. **(Fig. 11u, Supp data 6)**. Over-representation analysis of only 6 age DEGs specific to the male sex, showed enrichment of three processes represented in **Fig. 11t (Supp data 8)** but only with 3 total associated transcripts, *Tca3*, *Nrap*, and *Efhc2*. To determine the extent of overlap of age-related changes in each sex, Nrf2 WT and Nrf2 KO retinas, we intersected the lists of age DEGs in female Nrf2 WT and Nrf2 KO **(Supp Fig. 3c)** and in male Nrf2 WT and Nrf2 KO samples **(Supp Fig. 3d)** (from analyses **in Fig 12e-12p**). Venn representation intersecting the 49 age DEGs in Nrf2 WT and 69 age DEGs in Nrf2 KO female retina samples showed only 10 DEGs overlapped between genotypes while 39 and 59 age DEGs were specific for Nrf2 WT and Nrf2 KO genotypes, respectively **(Supp Fig. 3c, Supp data 9)**. Similarly, intersection of 41 age DEGs in Nrf2 WT and 33 age DEGs in Nrf2 KO male RPE-choroid showed only 7 DEGs shared across genotypes with 26 and 34 age DEGs being specific for Nrf2 WT and Nrf2 KO genotypes, respectively **(Supp Fig. 3d, Supp data 9)**. These comparisons support that modest differential expression changes in response to aging in the retina are influenced by Nrf2 expression status in both sexes.

## Discussion

Aging is the main risk factor implicated in the development of a variety of ocular degenerative/neurodegenerative diseases of the eye, including cataracts, glaucoma, diabetic retinopathy, and AMD^99,100^. Many reports support that NRF2 contributes to tissue homeostasis by activating protective pathways and upregulating endogenous antioxidants to neutralize excessive oxidative stress, as well as regulating metabolism, bioenergetics, and inflammation^17–24^. In the posterior segment of the eye, Nrf2 is expressed in neuroretina cell types, such as MGC, and RGCs, as well as in the RPE^25–27^ where dysregulation of its expression has been implicated in the pathogenesis of AMD. This mechanistic association with disease is supported by reduced NRF2 expression in the RPE of early AMD donors and with smoking, a major modifiable risk factor for AMD^28^, as well as by recent gene therapy studies in which AAV NRF2 rescued retinal structure and function in mouse models of oxidative stress^101^. While valuable knowledge on the role of oxidative stress in disease pathogenesis has been gained from preclinical studies that pair Nrf2 deficiency with injury models to mimic some aspects of AMD disease presentation^29–31,33,42^, how Nrf2 ablation interacts with normal aging, the main risk factor for AMD, to affect gene expression programs and the functions of retina and RPE cells is an outstanding, clinically relevant question. Our present studies expand on the observations of early AMD-like degeneration in Nrf2 null mice by Zhao and collaborators, which suggested increased vulnerability of the outer retina to age-related degeneration by disruption of Nrf2 signaling^32^. Here, we expand on these studies both at the functional and molecular levels by profiling visual acuity, dark-adapted ERG retina and RPE responses, and bulk-level transcriptomes from female and male mouse retina and RPE-choroid tissues at 3 months and 14 months of age. While scRNAseq studies or affinity purification approaches are important to focus on the cell-specific retinal, RPE, and choroidal cell populations^39,43,51,66–68,102^, our bulk-RNAseq studies, along with deconvolution analyses of cellular representation in the sequenced RNA, contribute the first comprehensive datasets characterizing the combined transcriptomic profiles of retina and RPE-choroid tissues with age, Nrf2 deletion, and sex as biological variables.

The OKT responses, measured in our experimental groups to assess how aging and genotype affect visual acuity, depend on a hierarchy of specialized cells and neural circuits, starting in the retina and extending through brainstem. This is distinguished from the ERG recordings that we performed here, which inform us about neural retina and RPE cells electrical activity in response to such variables, without the involvement of higher centers in the brain. Our analyses of visual function revealed a statistically significant, modest reduction of visual acuity with aging in both Nrf2 KO and Nrf2 WT mice that occurred in both male and female subjects, which was more accentuated in the Nrf2 KO group **(Fig. 2)**. Scotopic a-and b-wave analyses showed an age-dependent reduction in a- and b-wave amplitudes within each genotype group, maintained when analyzing females and males separately **(Figs. 3 and 4)**, indicating both rod photoreceptor and second-order neuron functional decline with age regardless of sex. Of note, we found a genotype effect at the young age, with Nrf2 KO having lower a- and b-wave amplitudes compared to their WT counterparts. These genotype differences converged later in life, at 14 mo, such that the amplitude differences were no longer observed. This loss of genotype effect on a- and b-wave responses at adult age could be explained by compensatory/adaptation mechanisms to overcome early functional deficits due to Nrf2 depletion, or by a “floor effect” in ERG responses reached with age alone that could not be further affected by genotype. The interaction effect of aging and Nrf2 expression status to reduce visual acuity was also observed in the analysis of dark-adapted electroretinographic c-wave responses, consistent with impaired RPE function **(Fig. 5)**. While our morphometry analyses could not discern superior from inferior hemispheres of the eye, the average measurements at equidistant points from the ON support no reduction in ONL thickness with Nrf2 deletion at the ages examined **(Fig. 6)**. Overall, functional and structural assessments are concordant with the idea that Nrf2 deletion impairs visual acuity and reduces of the amplitude of ERG responses in the retina and RPE at 14 mo without apparent photoreceptor cell death. Of note, a limitation of this study is that we did not focus on RPE/choroid morphology and structure, which would have potentially aided in the contextualization of c-wave ERG reduction and transcriptomic changes with age and Nrf2 deletion.

Earlier studies showed that the aging RPE (15 mo) expresses higher levels of the Nrf2 target genes *Nqo1*, *Gclm*, and *Hmox1* relative to the RPE of younger mice (2 mo) under unstressed conditions, suggesting an age-related increase in basal oxidative stress levels^31^. Moreover, the RPE of older mice exhibited impaired induction of the protective Nrf2 pathway in response to administration the oxidative stressor sodium iodate^31^. These observations are consistent with our RNAseq data comparing the expression of Nrf2 target genes between 14 mo and 3 mo RPE-choroid samples. As we only examine the effect of aging in the absence of any other stressors, we detected a modest age-related increase in the expression of Nrf2 target genes in the WT counterparts but not in the Nrf2 KO group **(Fig. 8c)**. Our neuroretina RNAseq profiling did not show a noticeable up-regulatory response of Nrf2 target genes **(Fig. 8e)** although these findings may have been skewed by the dominant abundance in neuroretina preparations of rod photoreceptor cells that could mask such anti-oxidant signatures in less abundant cell types including MGCs and RGCs **(Fig. 8e)**.

Our concerted experimental and analytical pipelines of RPE-choroid and retina expression profiles at the same ages, from the same cohort of animals, enhanced the relevance of the molecular findings of this study, supporting a timeframe of RPE-choroid responses to aging and genetic stress that likely precedes retinal degeneration. This notion is supported by a more pronounced differential expression response to aging and Nrf2 deletion in the RPE-choroid **(Fig 8. a-d)** than in the retina **(Fig 8. e-g)**. Our analytical approaches coupled with pathway analyses allowed us to uncover differential age and genotype effects on RPE-choroid gene expression. The intersection of age DEGs (14 mo vs 3 mo) in Nrf2 KO and Nrf2 WT RPE-choroid samples revealed 35% overlap between genotypes, as well as 26% and 5% of age DEGs being unique to the Nrf2 KO and Nrf2 WT groups, respectively. Canonical pathways with positive z-score values, represented by age-related gene expression changes across genotypes, were consistent with inflammatory, stress and disease mechanism/responses (neutrophil degranulation, phagosome formation, neuroinflammation, TLR receptor cascades, NK cell signaling, chemokines, cancer, MS, production of NO and ROS, NAFLD signaling) **(Fig. 9a)**. Genes with shared representation in these processes include TLR transcripts, *Tlr2*, *Tlr7*, *Tlr8*, *Tlr9,* and *Tlr13*. We previously reported mouse RPE, in response to aging, up-regulates *Tlr7* and *Tlr13* (mouse ortholog for human TLR3, 7, and 9)^43^, consistent with the first report identifying TLR gene expression (TLRs 1-7, 9, and 10) in human RPE cells, as part of a rapid response system to limit damage to non-regenerating retinal cells^103^. The findings of high expression of TLR2, TLR3 and TLR9 in human choroidal endothelial cells^104^ as well as enrichment of TLR2 and TLR7 in both macrophage (resident and infiltrating) and endothelial (artery, vein, and choriocapillaris) cell compartments of human choroid tissue by scRNAseq^105^ further support that the RPE-choroid environment elicits innate immune responses, which in our studies are up-regulated by age in both Nrf2 WT and Nrf2 KO mice.

Key characteristics of cytoskeletal fibers in the aged RPE/choroid include size reduction, decreased elongation, and collapsed morphology^106^. The structural integrity of the human RPE/choroid is largely dependent on the quality of the extracellular matrix, which is produced, organized, and maintained by fibroblasts that impact the RPE and choroid integrity and function^107^. In our study, canonical pathways with negative z-score values, represented by age-related gene expression changes in RPE-choroid across genotypes, were consistent with extracellular matrix remodeling processes. Transcripts with roles in collagen biosynthesis that are down-regulated with aging were *Col1a1 Col1a2, Col3a1, Col4a1, Col4a2, and Col5a1* **(Supp. data 3)**. This response was consistent with a reported negative correlation of expression of the supramolecular fiber organization pathway genes with age, in most cell types in human choroid tissue^107^ and is suggestive of age-related structural changes that are evident already in the adulthood.

An interesting finding of our study of age differences within genotypes was that ∼40% of age DEGs (539) were unique to Nrf2 KO RPE-choroid tissue, revealing an age-related signature that was absent in WT counterparts **(Fig. 9a)**. These unique age-related changes were associated with negative activation z-scores for fibrotic processes (i.e. pulmonary fibrosis, FAK, wound healing signaling), Notch/Hey1 signaling, and epithelial membrane protein signaling, while positive activation z-scores for three inflammation-related pathways. Activated neutrophils release nuclear and granular contents to form neutrophil extracellular traps (NETs), web-like DNA structures implicated in several neuroinflammatory conditions and a recent study reported that Nrf2 deficiency promotes brain blood barrier permeability, ICAM-1-mediated leukocyte transmigration, and neutrophil extracellular trap (NET) formation^108^. Our finding of altered expression of genes in the neutrophil extracellular trap signaling with aging specifically in Nrf2 KO RPE choroid **(Supp. data 3)** suggests acceleration of this process with aging in the absence Nrf2 signaling.

In the differential expression analysis of RPE-choroid but not of retina, we found a set of 184 DEGs with interaction effect of age and genotype. Pathway comparison analysis of the 184 DEGs revealed differential activation z-scores for several pathways, including complex IV assembly, respiratory electron transport and mitochondrial dysfunction across paired comparisons **(Supp. Fig 1)**. NRFs (NRF1 and NRF2) act on most nuclear genes that specify subunits of the five respiratory complexes in the mitochondrial inner membrane, of which complex IV facilitates the final transfer of electrons in the electron transport chain (ETC) from cytochrome c to molecular oxygen forming water and thus plays a key role in mitochondrial oxidative phosphorylation^109,110^. This provides a mechanism to maintain a balance between the metabolic rate of cells and the production of reactive oxygen species (ROS), which are produced as a byproduct of mitochondrial activity and can be deleterious if in excess (I.e., not sufficiently neutralized). Our analysis predicted inhibition of Complex IV assembly and respiratory electron transport in the Nrf2 KO vs Nrf2 WT at 3 mo as well as in the 14 mo vs 3 mo WT RPE-choroid comparisons. Conversely, activation of the same pathways was predicted for the Nrf2 KO vs Nrf2 WT at 14mo and for 14 mo vs 3mo Nrf2 KO RPE-choroid comparisons. The prediction for mitochondrial dysfunction was of activation for the Nrf2 KO vs Nrf2 WT at 3 mo as well as in the 14 mo vs 3 mo WT RPE-choroid comparisons and inhibition for the Nrf2 KO vs Nrf2 WT at 14 mo and for 14 mo vs 3 mo Nrf2 KO RPE-choroid comparisons.

Cigarette smoking (CS) is the strongest environmental risk factor for AMD and Nrf2 likely plays a protective role against this complex chemical oxidant^28^. CS has been shown to reprogram the transcriptome and epigenome of RPE cells *in vitro* and in preclinical models^70,111^, with findings correlated to human RPE from a smoker and early-stage AMD patient^111^. An earlier RNAseq study from Wang and collaborators^112^ showed that following chronic CS exposure, the RPE-choroid tissue exhibited dysregulated innate immune and morphogenic responses, consistent with an attempt to maintain cell differentiated states despite smoke-induced injury. In their recent snRNAseq work, Singh and collaborators^111^ reported up- and down-regulated DEGs in RPE cells from young mice intravitreally injected with cigarette smoke condensate (CSC). Under the assumption that smoking induces an accelerated aging response in RPE-choroid cells by increasing oxidative stress, and thereby exhausting Nrf2-mediated cytoprotective mechanisms, we contrasted our lists of age DEGs in the RPE-choroid of Nrf2 WT and Nrf2 KO mice **(Fig. 8a)** with reported DEGs in the RPE-choroid of mice chronically exposed to CS^112^ **(Supp Fig. 4, Supp data 10)**, as well as with DEGs in RPE cells in response to intravitreal CSC^111^ **(Supp Fig. 5, Supp data 10)**. The intersection of age DEGs in the RPE-choroid of Nrf2 WT and Nrf2 KO mice with a set of 2,012 DEGs induced by CS in mouse RPE-choroid (from Table S1 from^112^) rendered 230 DEGs shared across studies. From these, 96 were specifically common between the aged Nrf2 KO and CS-treated RPE-choroids **(Supp Fig. 4, Supp data 10)**. Some of these DEGs were over-represented in biological processes such as pigmentation, xenobiotic/amine transport, response to IL1β, and response to bacterial molecules, for the most part with discordant directionality of expression fold change across stress models.

Cross-referencing age DEGs from our study with a list of 408 (up-and down-regulated) DEGs specific to the RPE cell cluster in response to intravitreal CSC (combined from Tables S2-S4 from ^111^), we found that despite the very distinct nature of both studies, the intersection of these data sets revealed 14 DEGs across studies (all of which were DEGs in the Nrf2 KO 14 mo vs Nrf2 KO 3 mo comparison) **(Supp Fig. 5, Supp data 10)**. These shared

DEGs included C1qa, C1qb, C1qc among others, and were over-represented in humoral immune, TGF-β, and neuroinflammation responses. The results from these dataset intersections **(Supp Figs. 4-5)** suggest largely peculiar expression responses across experimental paradigms and that some specific age- and Nrf2 deletion-related changes in our bulk analyses could be contributed by RPE cells. Future studies, focusing specifically on aged RPE cells from Nrf2 deficient mice will enable more informative inter-study comparisons to elucidate mechanisms of RPE degeneration, as part of common or unique responses to different stressors.

In addition to analyzing the transcriptional profiles of RPE-choroid and neuroretina with aging and genotype, our studies considered sex as a biological variable, following NIH guidelines^113^, and based on published data of sex-divergences with aging in retina and RPE^43,88^. Our analyses revealed a small number of sex main effects (13-14) within each Nrf2 KO and Nrf2 WT RPE-choroid groups, mostly shared across ages and an interesting bias in the age main effects toward the female sex in both Nrf2 KO and Nrf2 WT RPE-choroid **(Fig. 11)**. Over-representation analysis of the female-specific age DEGs in the Nrf2 KO RPE-choroid enriched for growth factor (PDGF), immune, and ketone responses, and regulation of calcium. Homing of leukocytes, such as macrophages, to degeneration sites, is partly orchestrated by chemokines, such as CXCL1. CXCL1 transcript levels were reported to be up-regulated in macular and extramacular RPE-choroid of AMD donors^114^. In our study, *Cxcl1*, represented under regulation of cytosolic calcium concentration and leukocyte mediated immunity processes **(Fig 11, Supp. data 5)**, was differentially up-regulated with aging only in female Nrf2 KO RPE-choroid, suggesting a sex-specific predisposition toward activation of leukocyte homing responses in female tissue with normal aging. A gene represented in leukocyte immunity and production of molecular mediators of immune response processes was *Spon2*, coding for Spondin2. SPON2 is a secreted extracellular matrix protein, initially described as an essential pattern-recognition molecule in the initiation of the innate immune response^115^. In RNAseq studies of human RPE-choroid tissues, SPON2 expression was found to be enriched in fibroblasts, melanocytes, and NK cell clusters^116,117^. In our bulk-RNAseq of RPE-choroid, by deconvolution analysis, we found representation of transcriptomic signatures of stromal cell (largely composed of fibroblasts) and hematopoietic cell populations among others **(Fig. 7b)**. We also found *Spon2* was differentially down-regulated with age in Nrf2 KO RPE-choroid, specifically in females. Apart from a sex-specific response, this observation suggests that our mouse RNAseq analysis captures the transcriptome signature of cells putatively involved in the production of Spon2, which is translatable to the reported findings in the human RPE-choroid milieu.

As described for RPE-choroid, data analysis revealed a small number of sex main effects within each Nrf2 KO and Nrf2 WT retina groups, mostly shared across ages. We also observed a bias in the age main effects toward the female sex in both Nrf2 KO and Nrf2 WT retina **(Fig.12)**. PCA of expressed genes clearly separated retina transcriptomes by age in the first component in both genotype groups but a more defined segregation by sex in the second component was observed in the 3mo in the Nrf2 WT **(Fig. 12b)** compared to 3mo Nrf2 KO **(Fig. 12a)** retina transcriptomes.

Our comprehensive studies coupling visual function, retina structure, and gene expression profiles of the mouse retina and RPE-choroid allowed us to discover an interaction effect of aging and Nrf2 expression status to reduce visual acuity and to change bulk-tissue gene expression. Age and genotype effects were observed in both retina and RPE ERGs. A lower c-wave response was consistent with age and genotype effects on RPE-choroid gene expression. Of interest, our analyses support that with aging and higher susceptibility to oxidative stress, RPE and choroidal impairment precedes retina functional/structural decline. We provide functional data and transcriptomic datasets from the same cohort of mice that can be used to better understand mechanisms of a wide range of retinal degenerations. Our findings have clinical implications since aging and oxidative stress are risk factors for AMD, among other retinal diseases.

## Acknowledgements

The authors would like to thank Kendra Plafker and Adeline Machalinski for technical assistance, and Drs. Michael Elliott and John Ash for their critical comments during the preparation of the manuscript.

## Commercial relationships disclosures

B.L. Wells (none), A. Filiberti (none), D.R. Stanford (none), K.D. Pham (none), D. A. Walton (none), S.M. Plafker (none), W.M. Freeman (none), A.J. Chucair-Elliott (none)

## References

1. Pennington, K.L. & DeAngelis, M.M. Epidemiology of age-related macular degeneration (AMD): associations with cardiovascular disease phenotypes and lipid factors. Eye Vis (Lond) 3, 34 (2016).

2. Wong, W.L., et al. Global prevalence of age-related macular degeneration and disease burden projection for 2020 and 2040: a systematic review and meta-analysis. Lancet Glob Health 2, e106–116 (2014).

3. Brandl, C., et al. Features of Age-Related Macular Degeneration in the General Adults and Their Dependency on Age, Sex, and Smoking: Results from the German KORA Study. PLoS One 11, e0167181 (2016).

4. Mohana Devi, S., Mahalaxmi, I., Kaavya, J., Chinnkulandhai, V. & Balachandar, V. Does epigenetics have a role in age related macular degeneration and diabetic retinopathy? Genes Dis 8, 279–286 (2021).

5. Kar, D., et al. Choriocapillaris Impairment Is Associated With Delayed Rod-Mediated Dark Adaptation in Age-Related Macular Degeneration. Invest Ophthalmol Vis Sci 64, 41 (2023).

6. Orozco, L.D., et al. Integration of eQTL and a Single-Cell Atlas in the Human Eye Identifies Causal Genes for Age-Related Macular Degeneration. Cell Rep 30, 1246–1259 e1246 (2020).

7. Orozco, L.D., et al. A systems biology approach uncovers novel disease mechanisms in age-related macular degeneration. Cell Genomics, 100302 (2023).

8. Seddon, J.M., et al. Histopathological Insights Into Choroidal Vascular Loss in Clinically Documented Cases of Age-Related Macular Degeneration. JAMA Ophthalmol 134, 1272–1280 (2016).

9. Sohn, E.H., et al. Choriocapillaris Degeneration in Geographic Atrophy. Am J Pathol 189, 1473–1480 (2019).

10. Yu, D.Y. & Cringle, S.J. Oxygen distribution and consumption within the retina in vascularised and avascular retinas and in animal models of retinal disease. Prog Retin Eye Res 20, 175–208 (2001).

11. Strauss, O. The retinal pigment epithelium in visual function. Physiol Rev 85, 845–881 (2005).

12. Reichhart, N. & Strauss, O. Ion channels and transporters of the retinal pigment epithelium. Exp Eye Res 126, 27–37 (2014).

13. Strauss, O. The Retinal Pigment Epithelium. in Webvision: The Organization of the Retina and Visual System (eds. Kolb, H., Fernandez, E. & Nelson, R.) (Salt Lake City (UT), 1995).

14. Wimmers, S., Karl, M.O. & Strauss, O. Ion channels in the RPE. Prog Retin Eye Res 26, 263–301 (2007).

15. Kushwah, N., Bora, K., Maurya, M., Pavlovich, M.C. & Chen, J. Oxidative Stress and Antioxidants in Age-Related Macular Degeneration. Antioxidants (Basel) 12(2023).

16. Lambros, M.L. & Plafker, S.M. Oxidative Stress and the Nrf2 Anti-Oxidant Transcription Factor in Age-Related Macular Degeneration. Adv Exp Med Biol 854, 67–72 (2016).

17. Cano, M., et al. Nrf2 deficiency decreases NADPH from impaired IDH shuttle and pentose phosphate pathway in retinal pigmented epithelial cells to magnify oxidative stress-induced mitochondrial dysfunction. Aging Cell 20, e13444 (2021).

18. Wu, K.C., Cui, J.Y. & Klaassen, C.D. Beneficial role of Nrf2 in regulating NADPH generation and consumption. Toxicol Sci 123, 590–600 (2011).

19. Holmström, K.M., et al. Nrf2 impacts cellular bioenergetics by controlling substrate availability for mitochondrial respiration. Biol Open 2, 761–770 (2013).

20. Ludtmann, M.H., Angelova, P.R., Zhang, Y., Abramov, A.Y. & Dinkova-Kostova, A.T. Nrf2 affects the efficiency of mitochondrial fatty acid oxidation. Biochem J 457, 415–424 (2014).

21. Alsabaani, N.A., et al. Nrf-2-dependent antioxidant and anti-inflammatory effects underlie the protective effect of esculeoside A against retinal damage in streptozotocin-induced diabetic rats. Biomed Pharmacother 173, 116461 (2024).

22. Chen, X.L. & Kunsch, C. Induction of cytoprotective genes through Nrf2/antioxidant response element pathway: a new therapeutic approach for the treatment of inflammatory diseases. Curr Pharm Des 10, 879–891 (2004).

23. Li, N. & Nel, A.E. Role of the Nrf2-mediated signaling pathway as a negative regulator of inflammation: implications for the impact of particulate pollutants on asthma. Antioxid Redox Signal 8, 88–98 (2006).

24. Thimmulappa, R.K., et al. Nrf2 is a critical regulator of the innate immune response and survival during experimental sepsis. J Clin Invest 116, 984–995 (2006).

25. Datta, S., Cano, M., Ebrahimi, K., Wang, L. & Handa, J.T. The impact of oxidative stress and inflammation on RPE degeneration in non-neovascular AMD. Prog Retin Eye Res 60, 201–218 (2017).

26. Navneet, S., et al. Hyperhomocysteinemia-induced death of retinal ganglion cells: The role of Müller glial cells and NRF2. Redox Biol 24, 101199 (2019).

27. Zhang, J., et al. The Role of Nrf2/sMAF Signalling in Retina Ageing and Retinal Diseases. Biomedicines 11(2023).

28. Wang, L., et al. Nrf2 signaling modulates cigarette smoke-induced complement activation in retinal pigmented epithelial cells. Free Radic Biol Med 70, 155–166 (2014).

29. Wei, Y., et al. Nrf2 has a protective role against neuronal and capillary degeneration in retinal ischemia-reperfusion injury. Free Radic Biol Med 51, 216–224 (2011).

30. Larabee, C.M., et al. Loss of Nrf2 exacerbates the visual deficits and optic neuritis elicited by experimental autoimmune encephalomyelitis. Mol Vis 22, 1503–1513 (2016).

31. Sachdeva, M.M., Cano, M. & Handa, J.T. Nrf2 signaling is impaired in the aging RPE given an oxidative insult. Exp Eye Res 119, 111–114 (2014).

32. Zhao, Z., et al. Age-related retinopathy in NRF2-deficient mice. PLoS One 6, e19456 (2011).

33. Zhao, Z., et al. gammadelta T cells as a major source of IL-17 production during age-dependent RPE degeneration. Invest Ophthalmol Vis Sci 55, 6580–6589 (2014).

34. Rowan, S., et al. A low glycemic diet protects disease-prone Nrf2-deficient mice against age-related macular degeneration. Free Radic Biol Med 150, 75–86 (2020).

35. Wu, D.M., et al. Nrf2 overexpression rescues the RPE in mouse models of retinitis pigmentosa. *JCI Insight* 6(2021).

36. Chan, K., Lu, R., Chang, J.C. & Kan, Y.W. NRF2, a member of the NFE2 family of transcription factors, is not essential for murine erythropoiesis, growth, and development. Proc Natl Acad Sci U S A 93, 13943–13948 (1996).

37. Bowes, C., et al. Retinal degeneration in the rd mouse is caused by a defect in the β subunit of rod cGMP-phosphodiesterase. Nature 347, 677–680 (1990).

38. Sahu, B., et al. Presence of rd8 mutation does not alter the ocular phenotype of late-onset retinal degeneration mouse model. Mol Vis 21, 273–284 (2015).

39. Chucair-Elliott, A.J., et al. Translatomic response of retinal Muller glia to acute and chronic stress. Neurobiol Dis 175, 105931 (2022).

40. Chucair-Elliott, A.J., et al. Inducible cell-specific mouse models for paired epigenetic and transcriptomic studies of microglia and astroglia. Commun Biol 3, 693 (2020).

41. Prusky, G.T., Alam, N.M., Beekman, S. & Douglas, R.M. Rapid quantification of adult and developing mouse spatial vision using a virtual optomotor system. Invest Ophthalmol Vis Sci 45, 4611–4616 (2004).

42. Zyla K., W.D.A., Plafker K.S., Kovats S., Georgescu C., Brush R.S., Tytanic M., Agbaga M.P., Plafker S.M. Dietary protection against the visual and motor deficits induced by experimental autoimmune encephalomyelitis. Frontiers in Neurology in press(2023).

43. Chucair-Elliott, A.J., et al. Age- and sex-divergent translatomic responses of the mouse retinal pigmented epithelium. Neurobiol Aging 140, 41–59 (2024).

44. Cameron, M.A., Barnard, A.R. & Lucas, R.J. The electroretinogram as a method for studying circadian rhythms in the mammalian retina. Journal of Genetics 87, 459–466 (2008).

45. Liu, S., Yuan, F. & Xiang, M. Comparative analysis of electroretinogram with subdermal and invasive recording methods in mice. Biochemical and Biophysical Research Communications 655, 118–126 (2023).

46. Park, J.C., et al. Effects of normal aging on the mouse retina assessed by full-field flash and flicker electroretinography. Scientific Reports 13, 8860 (2023).

47. Robson, A.G., et al. ISCEV Standard for full-field clinical electroretinography (2022 update). Documenta Ophthalmologica 144, 165–177 (2022).

48. Chucair-Elliott, A.J., et al. Tamoxifen induction of Cre recombinase does not cause long-lasting or sexually divergent responses in the CNS epigenome or transcriptome: implications for the design of aging studies. Geroscience (2019).

49. Clough, E., et al. NCBI GEO: archive for gene expression and epigenomics data sets: 23-year update. Nucleic Acids Res 52, D138–d144 (2024).

50. Liao, Y., Wang, J., Jaehnig, E.J., Shi, Z. & Zhang, B. WebGestalt 2019: gene set analysis toolkit with revamped UIs and APIs. Nucleic Acids Res 47, W199–w205 (2019).

51. Lehmann, G.L., et al. Single-cell profiling reveals an endothelium-mediated immunomodulatory pathway in the eye choroid. J Exp Med 217(2020).

52. Li, J., et al. Comprehensive single-cell atlas of the mouse retina. iScience 27, 109916 (2024).

53. Newman, A.M., et al. Determining cell type abundance and expression from bulk tissues with digital cytometry. Nat Biotechnol 37, 773–782 (2019).

54. Rowan, S., et al. Aged Nrf2-Null Mice Develop All Major Types of Age-Related Cataracts. Invest Ophthalmol Vis Sci 62, 10 (2021).

55. Choudhary, M., Tayyari, F., Handa, J.T. & Malek, G. Characterization and identification of measurable endpoints in a mouse model featuring age-related retinal pathologies: a platform to test therapies. Lab Invest 102, 1132–1142 (2022).

56. Birch, D.G. & Anderson, J.L. Standardized Full-Field Electroretinography: Normal Values and Their Variation With Age. Archives of Ophthalmology 110, 1571–1576 (1992).

57. Ferdous, S., et al. Age-Related Retinal Changes in Wild-Type C57BL/6J Mice Between 2 and 32 Months. Investigative Ophthalmology & Visual Science 62, 9–9 (2021).

58. Gresh, J., Goletz, P.W., Crouch, R.K. & Rohrer, B. Structure–function analysis of rods and cones in juvenile, adult, and aged C57BL/6 and Balb/c mice. Visual Neuroscience 20, 211–220 (2003).

59. Kolesnikov, A.V., Fan, J., Crouch, R.K. & Kefalov, V.J. Age-Related Deterioration of Rod Vision in Mice. The Journal of Neuroscience 30, 11222–11231 (2010).

60. Peachey, N.S., Stanton, J.B. & Marmorstein, A.D. Noninvasive recording and response characteristics of the rat dc-electroretinogram. Visual Neuroscience 19, 693–701 (2002).

61. Wu, J., Peachey, N.S. & Marmorstein, A.D. Light-evoked responses of the mouse retinal pigment epithelium. J Neurophysiol 91, 1134–1142 (2004).

62. Wang, K., Zheng, M., Lester, K.L. & Han, Z. Light-induced Nrf2−/− mice as atrophic age-related macular degeneration model and treatment with nanoceria laden injectable hydrogel. Scientific Reports 9, 14573 (2019).

63. García-Llorca, A., Ólafsson, K.H., Sigurdsson, A.T. & Eysteinsson, T. Progressive Cone-Rod Dystrophy and RPE Dysfunction in Mitfmi/+ Mice. Genes 14, 1458 (2023).

64. Sugita, Y., Yamamoto, H., Maeda, Y. & Furukawa, T. Influence of Aging on the Retina and Visual Motion Processing for Optokinetic Responses in Mice. Front Neurosci 14, 586013 (2020).

65. Macosko, E.Z., et al. Highly Parallel Genome-wide Expression Profiling of Individual Cells Using Nanoliter Droplets. Cell 161, 1202–1214 (2015).

66. Pauly, D., et al. Cell-Type-Specific Complement Expression in the Healthy and Diseased Retina. Cell Rep 29, 2835–2848 e2834 (2019).

67. Heng, J.S., et al. Comprehensive analysis of a mouse model of spontaneous uveoretinitis using single-cell RNA sequencing. Proc Natl Acad Sci U S A (2019).

68. Hoang, T., et al. Gene regulatory networks controlling vertebrate retinal regeneration. Science 370(2020).

69. Lukowski, S.W., et al. A single-cell transcriptome atlas of the adult human retina. EMBO J 38, e100811 (2019).

70. Wang, J., et al. ATAC-Seq analysis reveals a widespread decrease of chromatin accessibility in age-related macular degeneration. Nat Commun 9, 1364 (2018).

71. Sun, L., et al. Single cell RNA sequencing (scRNA-Seq) deciphering pathological alterations in streptozotocin-induced diabetic retinas. Experimental Eye Research 210, 108718 (2021).

72. O’Koren, E.G., et al. Microglial Function Is Distinct in Different Anatomical Locations during Retinal Homeostasis and Degeneration. Immunity 50, 723–737.e727 (2019).

73. Reagan, A.M., et al. Age-related focal loss of contractile vascular smooth muscle cells in retinal arterioles is accelerated by caveolin-1 deficiency. Neurobiol Aging 71, 1–12 (2018).

74. Romay, M.C., et al. Age-related loss of Notch3 underlies brain vascular contractility deficiencies, glymphatic dysfunction, and neurodegeneration in mice. J Clin Invest 134(2024).

75. Dayalan Naidu, S., et al. CD5L is a target of transcription factor Nrf2. Biochemical and Biophysical Research Communications 776, 152225 (2025).

76. Takata, N., et al. An Eye Organoid Approach Identifies Six3 Suppression of R-spondin 2 as a Critical Step in Mouse Neuroretina Differentiation. Cell Reports 21, 1534–1549 (2017).

77. Yu, X., et al. Myelin regulatory factor deficiency is associated with the retinal photoreceptor defects in mice. Visual Neuroscience 38, E005 (2021).

78. Chowers, I., et al. The iron carrier transferrin is upregulated in retinas from patients with age-related macular degeneration. Invest Ophthalmol Vis Sci 47, 2135–2140 (2006).

79. Farkas, R.H., et al. Increased expression of iron-regulating genes in monkey and human glaucoma. Invest Ophthalmol Vis Sci 45, 1410–1417 (2004).

80. Coffey, P.J., et al. Complement factor H deficiency in aged mice causes retinal abnormalities and visual dysfunction. Proc Natl Acad Sci U S A 104, 16651–16656 (2007).

81. Ding, J.D., et al. Expression of human complement factor H prevents age-related macular degeneration-like retina damage and kidney abnormalities in aged Cfh knockout mice. Am J Pathol 185, 29–42 (2015).

82. Sun, X., Zhan, M., Sun, X., Liu, W. & Meng, X. C1GALT1 in health and disease. Oncol Lett 22, 589 (2021).

83. Adijanto, J., et al. The retinal pigment epithelium utilizes fatty acids for ketogenesis. J Biol Chem 289, 20570–20582 (2014).

84. Hunter, A., et al. DNA methylation is associated with altered gene expression in AMD. Invest Ophthalmol Vis Sci 53, 2089–2105 (2012).

85. Kim, H.J., Kim, H., Lee, J.H. & Hwangbo, C. Toll-like receptor 4 (TLR4): new insight immune and aging. Immun Ageing 20, 67 (2023).

86. Ozato, K., Tsujimura, H. & Tamura, T. Toll-like receptor signaling and regulation of cytokine gene expression in the immune system. Biotechniques **Suppl**, 66–68, 70, 72 passim (2002).

87. Tsioti, I., et al. Endothelial Toll-like receptor 4 is required for microglia activation in the murine retina after systemic lipopolysaccharide exposure. J Neuroinflammation 20, 25 (2023).

88. Du, M., et al. Retinal gene expression responses to aging are sexually divergent. Mol Vis 23, 707–717 (2017).

89. Qin, Y.J., et al. Autonomic nervous system imbalance in diabetic mouse choroids. Tissue Cell 94, 102798 (2025).

90. Kärkkäinen, V., et al. Neuroinflammation Markers in Tear Fluid of Mild Alzheimer’s Disease. J Mol Neurosci 75, 73 (2025).

91. Sack, G.H., Jr. Serum amyloid A - a review. Mol Med 24, 46 (2018).

92. Moore, Y.F., Lambracht-Washington, D., Tabaczewski, P. & Fischer Lindahl, K. Murine MHC class Ib gene, H2-M2, encodes a conserved surface-expressed glycoprotein. Immunogenetics 56, 1–11 (2004).

93. Etebar, F., Whatmore, P., Harkin, D.G. & Dando, S.J. Tissue-Specific Immune Transcriptional Signatures in the Bordering Tissues of the Mouse Retina and Brain. Invest Ophthalmol Vis Sci 65, 42 (2024).

94. Xie, W., Gao, L., Deng, H., Liu, D. & Pang, Q. The Evolution, Oligomerization, Function, and Action Mechanism of α2-Macroglobulin. Cells 15(2026).

95. Schmalen, A., et al. Proteomic Phenotyping of Stimulated Muller Cells Uncovers Profound Pro-Inflammatory Signaling and Antigen-Presenting Capacity. Front Pharmacol 12, 771571 (2021).

96. Mangold, C.A., et al. CNS-wide Sexually Dimorphic Induction of the Major Histocompatibility Complex 1 Pathway With Aging. J Gerontol A Biol Sci Med Sci 72, 16–29 (2017).

97. Bhutto, I.A., McLeod, D.S., Merges, C., Hasegawa, T. & Lutty, G.A. Localisation of SDF-1 and its receptor CXCR4 in retina and choroid of aged human eyes and in eyes with age related macular degeneration. Br J Ophthalmol 90, 906–910 (2006).

98. Elamaa, H., et al. Angiopoietin-4-dependent venous maturation and fluid drainage in the peripheral retina. Elife 7(2018).

99. Klein, R. & Klein, B.E. The prevalence of age-related eye diseases and visual impairment in aging: current estimates. Invest Ophthalmol Vis Sci 54, ORSF5–ORSF13 (2013).

100. Luu, J. & Palczewski, K. Human aging and disease: Lessons from age-related macular degeneration. Proc Natl Acad Sci U S A 115, 2866–2872 (2018).

101. Gardner, A., et al. AAV NRF2 gene therapy preserves retinal structure and function in rodent models of oxidative damage. Mol Ther 34, 3245–3269 (2026).

102. Chucair-Elliott, A.J., et al. Comparative analysis of In vivo endothelial cell translatomes across central nervous system vascular beds. Exp Eye Res 248, 110101 (2024).

103. Kumar, M.V., Nagineni, C.N., Chin, M.S., Hooks, J.J. & Detrick, B. Innate immunity in the retina: Toll-like receptor (TLR) signaling in human retinal pigment epithelial cells. J Neuroimmunol 153, 7–15 (2004).

104. Stewart, E.A., Wei, R., Branch, M.J., Sidney, L.E. & Amoaku, W.M. Expression of Toll-like receptors in human retinal and choroidal vascular endothelial cells. Experimental Eye Research 138, 114–123 (2015).

105. Voigt, A.P., et al. Choroidal endothelial and macrophage gene expression in atrophic and neovascular macular degeneration. Hum Mol Genet 31, 2406–2423 (2022).

106. Ramrattan, R.S., et al. Morphometric analysis of Bruch’s membrane, the choriocapillaris, and the choroid in aging. Invest Ophthalmol Vis Sci 35, 2857–2864 (1994).

107. Huang, L., et al. Dynamic human retinal pigment epithelium (RPE) and choroid architecture based on single-cell transcriptomic landscape analysis. Genes Dis 10, 2540–2556 (2023).

108. Muneer, P.M.A., Bhowmick, S., Poovanthodi, Y.A. & Alikunju, S. Nrf2 regulates ICAM-1-mediated neutrophil extracellular trap formation after traumatic brain injury. bioRxiv (2026).

109. Pulliam, D.A., et al. Complex IV-deficient Surf1(-/-) mice initiate mitochondrial stress responses. Biochem J 462, 359–371 (2014).

110. Scarpulla, R.C. Nuclear control of respiratory chain expression by nuclear respiratory factors and PGC-1-related coactivator. Ann N Y Acad Sci 1147, 321–334 (2008).

111. Singh, K.K., et al. Molecular underpinnings of induced degenerative heterogeneity in the retinal pigment epithelium. Proc Natl Acad Sci U S A 123, e2505412123 (2026).

112. Wang, L., et al. Retinal pigment epithelium transcriptome analysis in chronic smoking reveals a suppressed innate immune response and activation of differentiation pathways. Free Radic Biol Med 156, 176–189 (2020).

113. Clayton, J.A. & Collins, F.S. Policy: NIH to balance sex in cell and animal studies. Nature 509, 282–283 (2014).

114. Newman, A.M., et al. Systems-level analysis of age-related macular degeneration reveals global biomarkers and phenotype-specific functional networks. Genome Med 4, 16 (2012).

115. He, Y.W., et al. The extracellular matrix protein mindin is a pattern-recognition molecule for microbial pathogens. Nat Immunol 5, 88–97 (2004).

116. Voigt, A.P., et al. Single-cell transcriptomics of the human retinal pigment epithelium and choroid in health and macular degeneration. Proc Natl Acad Sci U S A 116, 24100–24107 (2019).

117. Collin, J., et al. Single-cell RNA sequencing reveals transcriptional changes of human choroidal and retinal pigment epithelium cells during fetal development, in healthy adult and intermediate age-related macular degeneration. Hum Mol Genet 32, 1698–1710 (2023).

