## Supplemental Figures for "Functional and molecular profiles of the neuroretina and retinal pigmented epithelium-choroid in response to aging and Nrf2 deletion"

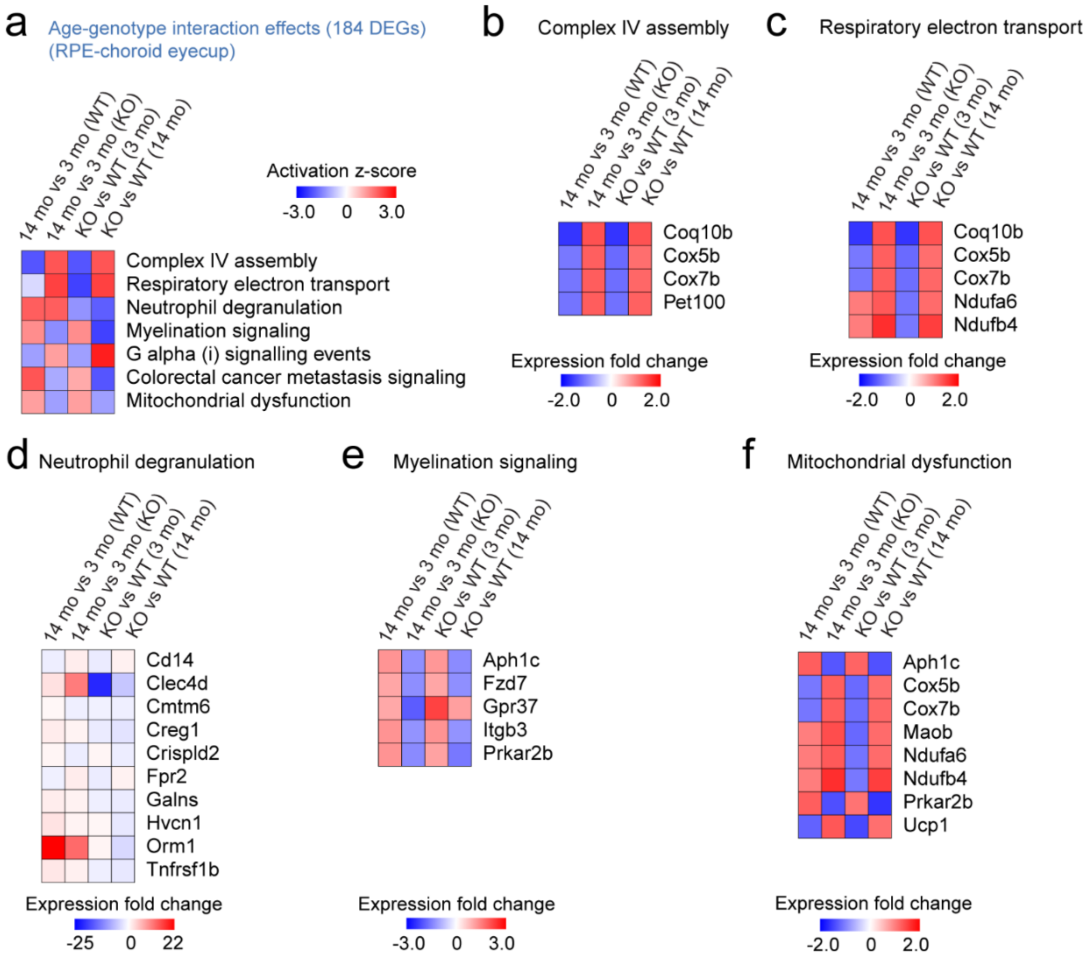


**Supplemental Figure 1. Comparison of RPE-choroid profile with aging and Nrf2 expression status reveals age-genotype interaction effects.** Pathway analysis revealed that the 184 age-genotype DEGs identified by 2-way ANOVA (Fig 8a and Fig 8d) were implicated in mitochondrial physiology (complex IV assembly, respiratory electron transport, mitochondrial dysfunction), as well as specific components if neutrophil degranulation and myelination signaling. **a:** Heatmap depicts predicted activation scores for all age and genotype comparisons. **b-f:** Heatmaps indicate expression fold change for genes in the pathways in (a).

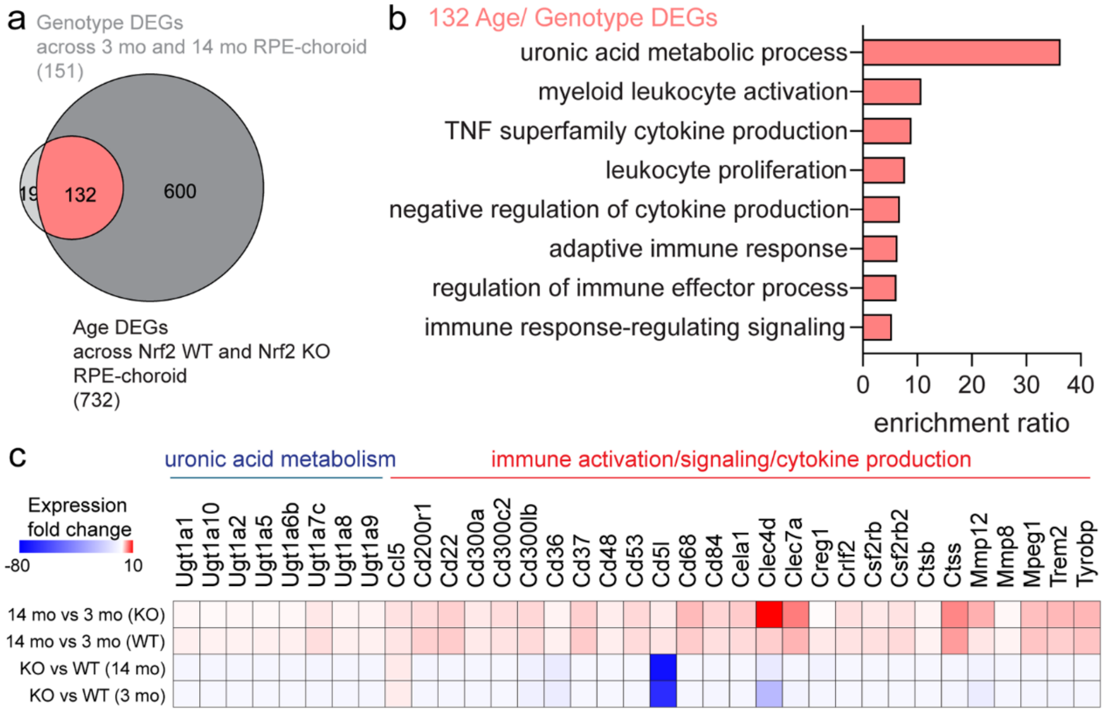


**Supplemental Figure 2. Contextualization of the differential gene expression profile of the RPE-choroid in response to aging and Nrf2 deletion. (con't). a:** The intersection between genotype DEGs common to 3mo and 14 mo RPE-choroid and age DEGs common to Nrf2 WT and Nrf2 KO RPE-choroid shows 132 genes, of which ORA via Webgestalt is shown in **b**. **c:** Heatmap represents expression fold change for genes represented in processes in **b**, for the Nrf2 KO vs Nrf2 WT (3mo), Nrf2 KO vs Nrf2 WT (14 mo), 14 mo vs 3 mo (Nrf2 KO), and 14 mo vs 3 mo (Nrf2 WT) RPE-choroid comparisons.

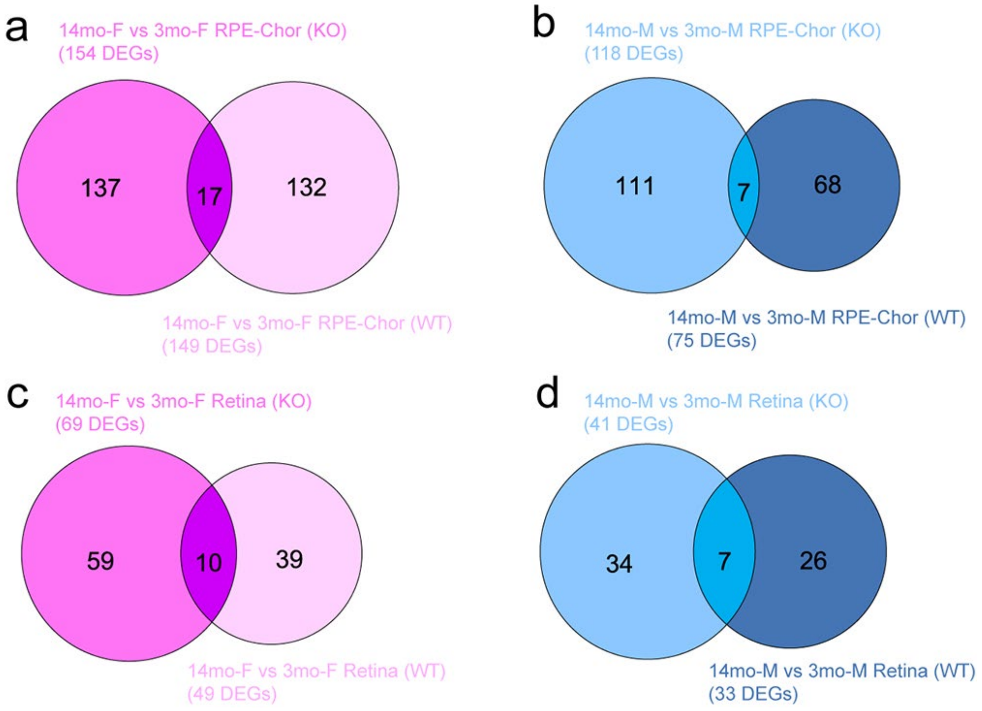


**Supplemental Figure 3. Age-related changes in female and male RPE-choroid and retina across genotypes.**

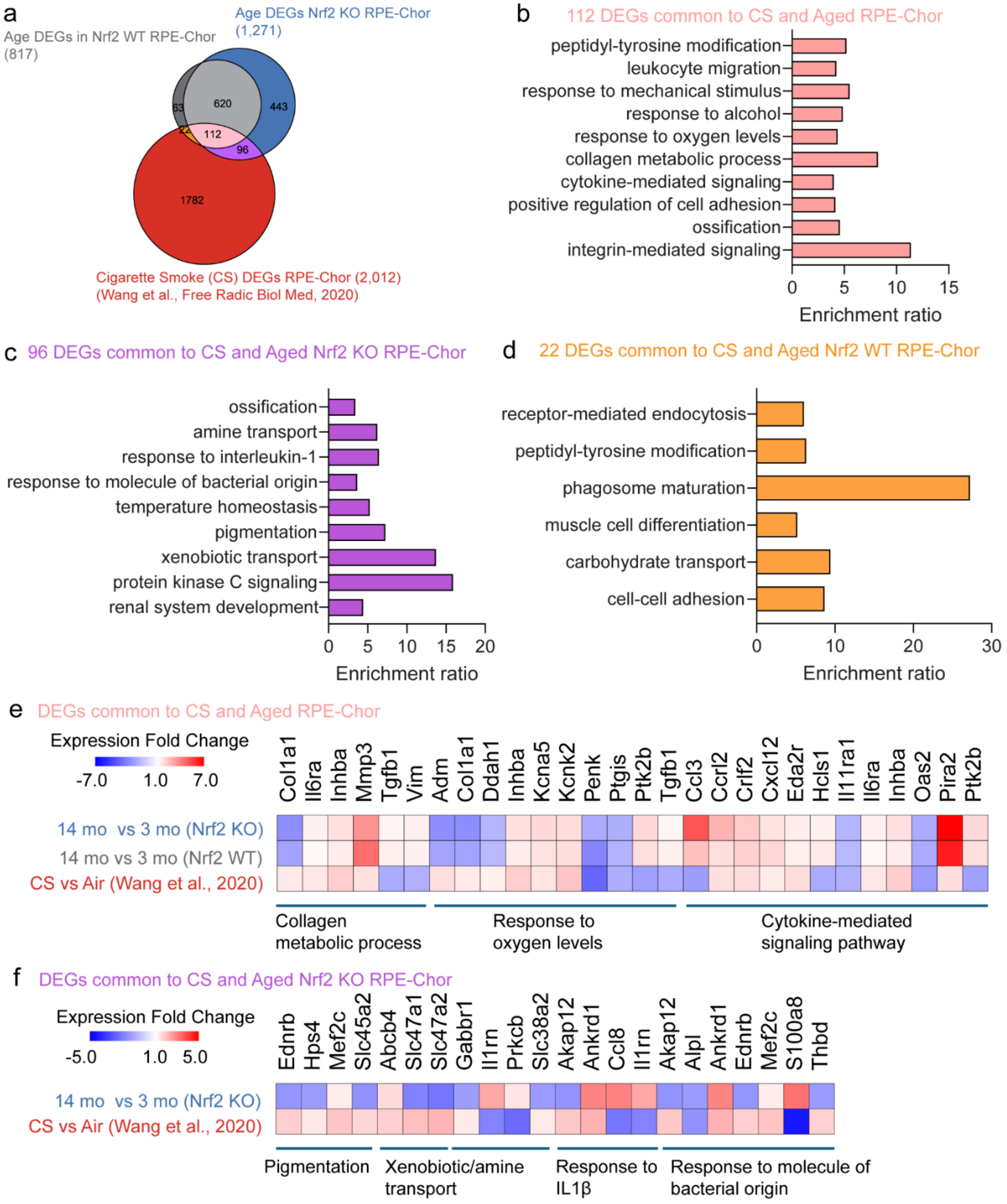


**Supplemental Figure 4. Age-related changes in changes in the RPE transcriptomic response to chronic smoking. a:** A list of 2,012 DEGs in RPE-choroid following 6 months of exposure to cigarette smoking (CS) (Wang et al., 2020) was intersected with sets of age DEGs in RPE-choroid from Nrf2 WT and Nrf2 KO mice, from this study. **b-d:** Over-representation analyses were conducted on subsets of DEGs shared between the three gene lists. **e-f:** Heatmaps illustrate expression fold change for select DEGs from analyses in b and c.

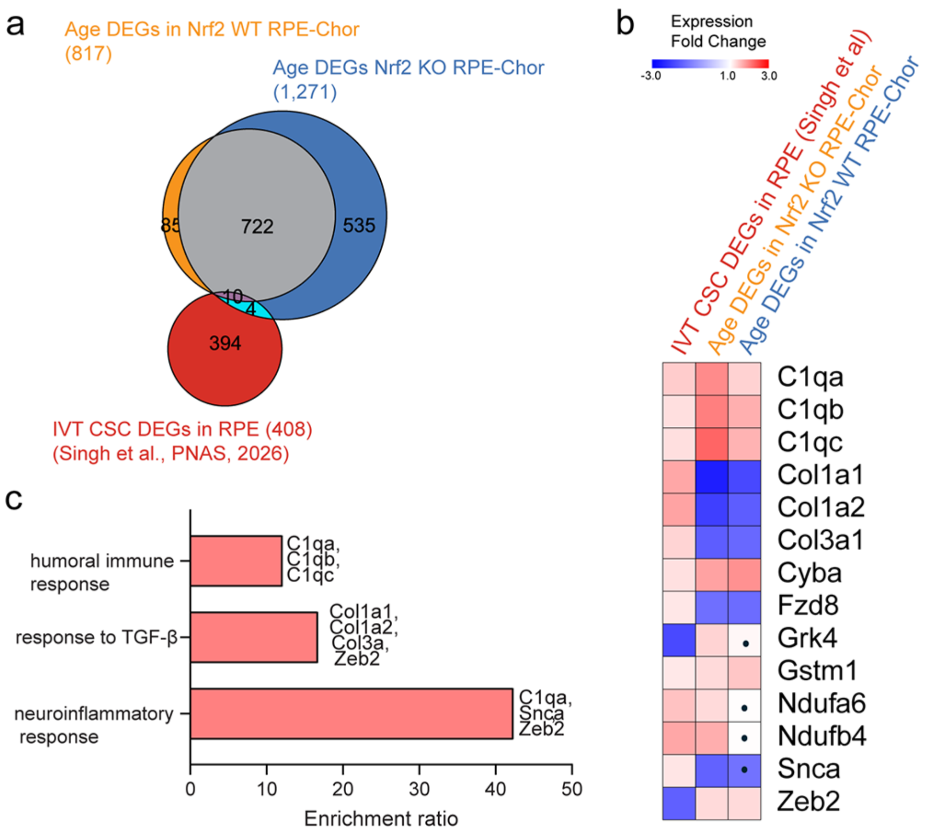


**Supplemental Figure 5. Age-related changes in the RPE-choroid in response to aging and Nrf2 deletion vs RPE cell transcriptomic responses to cigarette smoke condensate (CSD). a:** A list of 408 DEGs in the RPE cell cluster following 10 days of intravitreal cigarette smoke condensate (CSC) (Singh et al., 2026) was intersected with sets of age DEGs in RPE-choroid from Nrf2 WT and Nrf2 KO mice, from this study. **b:** Heatmap illustrates expression fold change for the 14 DEGs shared between CSC and our aging studies. **c:** Over-representation analysis was conducted on the 14 DEGs shared across stress models. Black dotes denote no statistically significant differential expression.
